# Affiliative behavioural phenotype and microglial activation predict social decision-making following immune activation

**DOI:** 10.1101/2025.11.23.690017

**Authors:** Emma R Hammond, Patrick K Monari, Elsa M Luebke, Aimee K Johnson, Anthony P Auger, Catherine A Marler

**Affiliations:** University of Wisconsin-Madison, Department of Psychology, Madison, WI, USA; University of Wisconsin-Madison, Neuroscience Training Program, Madison, WI, USA

**Keywords:** Inflammation, social decision-making, prefrontal cortex, microglia, lipopolysaccharide, affiliation, ultrasonic vocalizations

## Abstract

Social animals are faced with a critical decision when sick: to affiliate or withdraw. The behavioural response during an immune challenge varies by individual and species. However, the social and biological factors leading to variation in this decision during an immune challenge are unknown. Here, we explore the affiliative behavioural phenotype, “huddlers” and “non-huddlers,” as a social predictor, and microglial activation and peripheral cytokine expression as biological predictors of social-decision making during an immune challenge. To measure social-decision making, we recorded social approach and time spent in proximity to a familiar same-sex conspecific versus a novel object in the social California mouse following lipopolysaccharide (LPS) treatment. Behavioural phenotype predicted both the social response to LPS and the relationship between cytokine activity and social behaviour, suggesting that social experience regulates social decision-making during sick versus healthy conditions. Additionally, we found that elevated microglia activity in the dentate gyrus (DG) and medial prefrontal cortex (mPFC) negatively correlated with social behaviour, positioning these regions for future exploration for the role of microglia on social decision-making. Our results identify the predictive power of behavioural phenotypes on social response to sickness and a link between microglia and decision-making during sickness.

## 1. Introduction

While recovering from illness, animals adjust behaviour to adapt to different immune states. Although sickness behaviour is widespread in vertebrates [1], within species variation indicates that individuals are likely making social decisions to approach or withdraw based on internal state and social environment [2–4]. During an immune challenge, an individual may increase contact with an affiliative group to protect against predators and enhance the likelihood of recovery [3,5,6]; however, increased social contact may increase vulnerability to conspecific aggression and risk of disease transmission [7]. Alternatively, benefits of social withdrawal may include energy conservation, avoiding conspecific aggression, and protecting against the spread of disease [8,9]. Nevertheless, the social and biological bases for the decision to approach versus withdraw is unknown.

One crucial component for driving social-decision making is individual variation in behavioural phenotype and social relationships. Distinct behavioural phenotypes that predict differential responses to stress, social cues, and opioid release have been found in house mice, California mice, and starlings [10–12], emphasizing the importance of understanding individual variation within-species. Such variation may contribute to different responses during an immune challenge, especially when distinct social phenotype may alter social relationships. Rats, rhesus macaques, and house finches increase sociality to familiar conspecifics during an immune challenge or infection [3,4,13], a striking difference from the classical social withdrawal associated with sickness behavior [9]. Plasticity in social response to immune challenge is illustrated by research showing approach to familiar but avoidance of unfamiliar conspecific rats [4], but it is unknown if individual variation among familiar conspecifics relates to social relationship quality.

The monogamous California mouse (*Peromyscus californicus*) is a potent model for studying social decision making across immune states and varying social relationships. California mice express high variation in affiliation between same-sex bonds [14], allowing us to examine whether affiliation levels predict behavioural changes during an immune challenge. California mice form strong social bonds but vary in levels of huddling and affiliative vocalizations [15,16]. We therefore chose to examine whether individual variation in behavioural phenotype positively associates with increased affiliation during an immune challenge as tested through social decision-making and affiliative behaviours. Thus, we predicted that mice with stronger affiliative bonds would increase sociability during a mild LPS immune challenge.

The biological mechanisms driving social withdrawal during an immune response are well-established [1], but little is known about the biological mechanisms driving affiliation during an immune challenge. Here, we manipulated oxytocin (OT) and measured microglia activation as potential mechanisms driving social decisions during an immune response. The neuropeptide OT is a crucial regulator of social behaviour [17], can increase or decrease affiliation [18], and can mediate behavioral coordination in California mice [10,19]. OT neurons are also critical for immune function, and release OT in response to immune challenges [20]. Although OT effects on immune function are not fully understood, OT typically displays anti-inflammatory qualities [21]. Microglia, the primary immune cells of the brain, connect OT and neural and immune function [22]. Microglia are enriched with OT receptors, and OT can blunt inflammation by acting on microglia following an immune challenge [23]. We therefore predicted that OT would increase sociability and decrease microglia activation following an inflammatory challenge.

Microglia support cognitive processes such as cognitive flexibility, stress resilience, and social interactions [24,25]. Overactivity of microglia is associated with social impairments [26], whereas inhibiting microglia activity using minocycline is associated with increased sociability [27]. Research in humans demonstrates that minocycline alters decision-making style [28]. However, minocycline treatment is not region specific, making our understanding of how microglia may function in region-specific ways limited.

Although many brain regions influence the expression of complex social behaviour, we focused on three regions of interest: the dorsal dentate gyrus (DG), medial prefrontal cortex (mPFC), and medial amygdala (MeA). The dorsal DG is an important regulator of mood and affiliation versus withdrawal [29]. The infralimbic mPFC impacts avoidance and reward seeking behaviour [30], as well as cognitive flexibility necessary for altering social decision making under diverse contexts [31] such as internal immune state changes. The third brain region was the MeA, necessary for the organization of social behaviour through processing and differentiating social cues and mediating behavioural output [32]. To understand decisions to socially withdraw or approach in an inflammatory state, we must understand integration of social cues.

Despite the established effects of sickness on behaviour, the social and biological mechanisms driving social decision making during sickness are unknown. We predicted that 1) LPS would increase same-sex affiliation, 2) this increase would be associated with a prior affiliative relationship with the same-sex conspecific, 3) an OT antagonist (OTA) would block this increase and OT would elevate it, 4) elevated microglia activity in brain regions relevant to social behaviour would be associated with reduced affiliation following LPS treatment.

## 2. Methods

### (a) Animals

49 (24 male, 25 female) 3-6 month old California mice from an established colony at the University of Wisconsin-Madison were housed with a same-sex cagemate and maintained in accordance with the NIH Guide for the care and use of laboratory mice. Cagemates were siblings and unrelated same-sex individuals housed together since weaning, balanced across groups. One mouse per cage was randomly selected as the focal mouse; the familiar cagemate served as the social stimulus. Mice were housed in standard cages (48 × 27 × 16 cm) with aspen bedding, a nestlet, Purina 5015™ chow, and water *ad libitum*. Housing conditions were 20 - 23° C on a 14L:10D cycle (lights off at 9:00am CST). Behaviour testing occurred 1-3 hrs after lights-off. Behaviour was measured in 1) the sociability test (Methods 2.d) and 2) undisturbed observations (Methods 2.e), and scored by one observer naive to treatment and sex.

### (b) Oxytocin and Oxytocin Antagonist Treatment

Mice were randomly assigned to control (n = 16), OT (n = 16), or OTA (n = 16) groups and received three intranasal (IN) treatments and three intraperitoneal (IP) injections on Days 2, 5, and 8. Controls received IN .9% saline (.4ul/g) and IP .9% saline (1ul/g). The OT group received IP saline and IN OT (Bachem, Torrance, CA, Prod #: 4016373) dissolved in .9% sterile saline at .5 IU/kg (.4ul/g), a dose that alters social behaviour in California mice [33] and equivalent to weight-adjusted human doses [34]. The OTA group received IN saline and 1mg/kg IP OTA (L-368,899) (Sigma-Aldrich, St. Louis, MO, Prod #: L2540) in .9% saline, a dose that alters social approach in California mice [35] and crosses the blood-brain barrier [36].

### (c) LPS Treatment

Lyophilized LPS (Sigma-Aldrich, St. Louis, MO, Prod #:L2762-10MG) from *E.coli* was dissolved in .9% saline. A .1mg/kg dose was selected based on prior work in *Peromyscus* [37]. This dose did not alter activity levels 24 hrs post-injection, as measured by time spent inactive compared to saline (p = .504) (Supplemental Figure 6.A). We used a low dose because higher doses suppress both social and non-social activity, interfering with social-decision making [38]. In contrast, low-grade inflammation disrupts social behaviour without affecting general activity [38], enabling isolation of inflammation-specific social effects. Overall weight decreased across the study (p <.001) with no differences between OT, OTA, and control groups (see Supplemental Figure 6.B). All mice received .9% saline (.4ul/g) on Day 1 and LPS on Day 4 to control for injection effects.

### (d) Sociability Test

The sociability test used a within-subject design. Test 1 occurred on Day 2, 24 hrs after saline (Figure 1), and Test 2 on Day 5, 24 hrs after LPS. Testing began 10 min after IN treatment and 30-min after IP injection to align with known effects of OT and OTA [10,35] and pharmacokinetic data [39,40]. Focal mice were habituated to the chamber for 20-min, 24 hrs before each test. During testing, the familiar cagemate was placed behind a mesh barrier in one side chamber and a novel object in the other [41]. The novel object (small saline bottle or binder clip, counterbalanced across tests) was chosen because of similarity in size to California mice. Behaviour was videotaped from a side-facing camera (Kicteck 4K, 60FPS) under red-light. Sociability was quantified as the ratio of time spent in the social versus object chamber [41].

**Figure 1.**
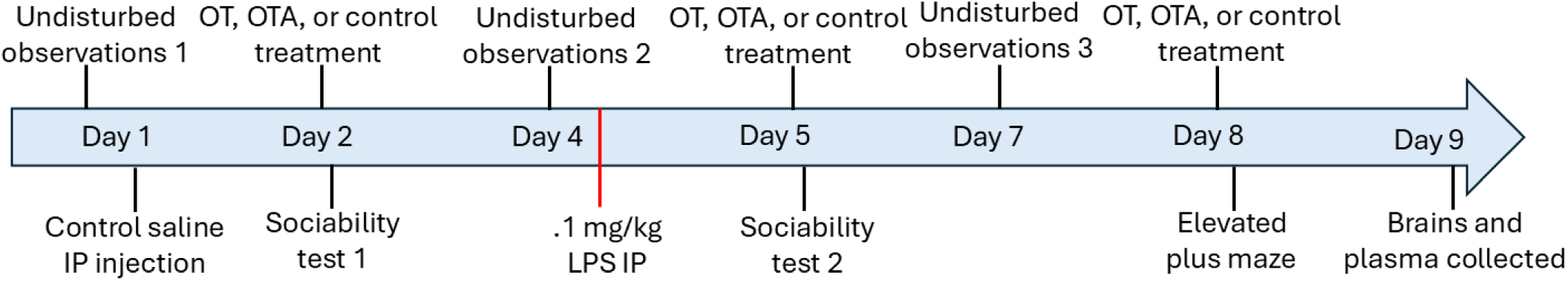
Timeline of methods. “Undisturbed observation 1” is also referred to as “baseline.”

### (e) Undisturbed Observations

Behaviours were recorded in the homecage on Day 1 (baseline), Day 2 (post OT/OTA/control), and Day 7 (two days after LPS, and one day after a second dose of OT/OTA/control), allowing comparison of baseline, saline + treatment, and LPS + treatment. Behaviour was videotaped from above (Kicteck 4K, 60FPS) under red-light and ultrasonic vocalizations (USVs) were recorded for 10 min. Food and water were removed during testing to prevent visual obstruction. Videos were scored in BORIS [42]. Proximity was defined as the two mice being within one mouse-length for > 5s and was measured as a cumulative duration over 10 min. All other behaviour definitions are described in Supplemental Table 1.

### (f) Huddling Analysis

Huddling was defined as flank-to-flank contact for >5 s. Because durations were skewed and zero-inflated (Supplemental Figure 1), data were binarized [43]. Mice were categorized as “huddlers” (n = 18) or “non-huddlers” (n = 31). “Huddlers” refers to mice that huddled during at least one of the three 10-min undisturbed observation periods; “non-huddlers” refers to mice that were never observed huddling (for details, see Supplemental Figure 1). This classification captured behavioural variation and is consistent with analyses using low/non-huddlers as a model of reduced social motivation in non-human primates [44]. Huddling data were missing from one of three time points for five mice due to file corruption or camera failure; these mice were excluded, yielding a final sample size of 44 for huddling analyses (23 females, 21 males).

### (g) Microglia Immunocytochemistry

Immunocytochemistry followed established protocols [33,45]. Images were collected from the infralimbic mPFC, dorsal DG, and MeA (Supplemental Figure 2). For each region, three images were acquired on a Zeiss 710 confocal microscope using a 20x objective. 10 microglia per region were randomly selected and scored on a 0-5 activation scale based on primary process number, number of processes with branches, and CD68 inclusions [46], details in Supplemental Methods 8.d.

### (h) Electrochemiluminescence Immunoassay

Inflammatory cytokine concentrations were quantified using a Meso Scale Discovery’s V-Plex Proinflammatory Panel 1 Mouse Kit (Meso Scale Discovery, Rockville, MD, USA, Cat. #K15048D), following manufacturer instructions and prior publications [47]. This multiplex electrochemiluminescent immunoassay measured serum concentrations of IFN-γ, IL-1ꞵ, IL-2, IL-4, IL-5, IL-6, IL-10, IL-12p70, KC/GRO and TNF-ɑ. IL-4 was excluded due to undetectable levels (Supplemental Methods 8.e). The assay runs 40 samples at a time, thus, 40 samples were analyzed (22 males and 18 females: control n = 15, OT n = 14, OTA n = 11). Sex was included as a covariate in cytokine analyses, but sex differences were not tested statistically due to limited power. A principal component analysis (PCA) was performed, consistent with previous cytokine studies [48,49]. PC1 loaded primarily on IL-6, IL-2, IL-12, and TNF-ɑ (Supplemental Figure 4), all mediators of the Th1 proinflammatory response [50,51]. Thus, PC1 was interpreted as proinflammatory cytokine activity.

### (i) Vocalization Analysis

USVs were recorded during each 10-min undisturbed observation using a Knowles FG ultrasonic microphone. Vocalizations were visually identified from spectrograms in Avisoft SASlab pro (Avisoft Bioacoustics, Berlin, Germany). DeepSqueak [52] was used to detect three call types California mouse call types: sweeps, barks, and sustained vocalizations (SVs) [10,15,16]. A post hoc denoising network was trained to reduce the false detection rate and improve detection accuracy. Sweeps occurred in all cages, but SVs and barks were rare and therefore excluded. For sweeps, call features were extracted using DeepSqueak. Call features included call length, principal frequency, slope, sinuosity, and power [53]. Because calls from individuals within a cage cannot be distinguished, vocalizations were analyzed per mouse pair.

### (j) Statistical Analyses

This study used a within-subjects design. Behavioural data were analyzed using linear mixed effects models (R version 4.1.3, RStudio), with Animal ID as a random effect. Data are reported as the mean ± SEM (“b” indicating mean), with significance set at p < .05. For undisturbed observations, baseline data (Day 1) were included as covariates when comparing effects of LPS + OT/OTA/control (Day 7) relative to saline + OT/OTA/control (Day 3). OT and OTA were treated as categorical variables and dummy-coded to compare differences relative to controls. Between-subjects data (e.g. cytokine data and microglia activation) were analyzed using linear regression. Effect sizes (R^2^) were interpreted as low ∼ .02, medium ∼ .13 and high ∼ .23 or greater [54]. When appropriate (Results 3.b; Supplemental Results 9.c), p-values were adjusted for multiple comparisons using Holm-Bonferonni [55]. Adjusted p-values are displayed. Model assumptions (normality, linearity, and homoscedasticity) were evaluated using case analysis.

Skewed data were log transformed as required. Cook’s distance was used to identify high-influence observations. If removal of influential points did not change the significance of interpretation (p < .05), all data were retained; if removal altered significance, results were reported as non-significant trends. Sex was included as a biological variable in all models. For clarity, sex-specific results are not shown in all figures but are reported in Supplemental Figure 15.

## 3. Results

### (a) Non-huddling male mice increased sociability following LPS

LPS increased sociability in non-huddler but not huddler males. We ran a mixed effects regression model including huddling, sex, OT/OTA/control treatment as between-subject variables, and LPS/saline as a within-subject variable, and sociability as the outcome variable. There was a three-way interaction between huddling, sex, and LPS, such that LPS influenced sociability in non-huddler males in the opposite direction of huddler males (b = .347 ± .163, p = .040, F(1,52) = 4.44, conditional R^2^ = .35, marginal R^2^ = .13) (Figure 2). Among non-huddlers, the interaction between LPS and sex revealed that LPS increased sociability overall in males relative to females (b = .178 ± .084, p = .040, F(1,44) = 4.46, conditional R^2^ = .35, marginal R^2^ = .13). Effects in males were further dependent on behavioral phenotype. In non-huddler males, LPS increased sociability relative to after saline treatment (b = .168 ± .060, p = .008, F(1,46) = 7.74, conditional R^2^ = .35, marginal R^2^ = .13). There was no main effect of LPS on sociability in huddler males (p = .176) or on females (p = .874). There was also no effect of OT (p = .83) or OTA (p = .656) on sociability in males or females.

**Figure 2.**
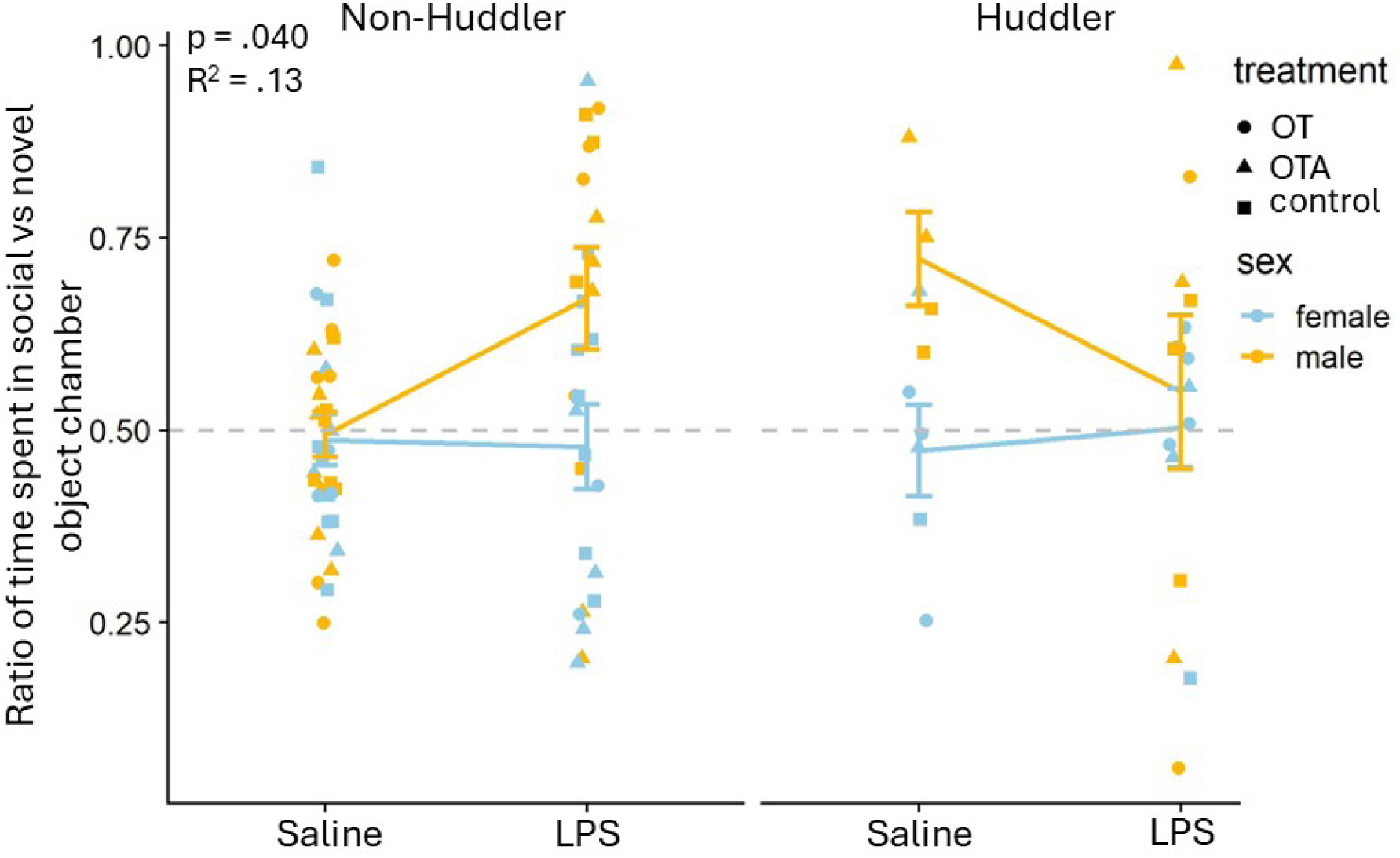
There was a three-way interaction between huddling, sex, and LPS, such that LPS influenced social preference in non-huddler males in the opposite direction as huddler males (p = .040, conditional R^2^ = .35, marginal R^2^ = .13). The gray line indicates equal time spent in the novel object and social chamber. Among non-huddlers, LPS increased social preference in males relative to females (p = .040, conditional R^2^ = .35, marginal R^2^ = .13). In non-huddler males, LPS increased the ratio of time spent relative to saline (p = .008, conditional R^2^ = .35, marginal R^2^ = .13).

### (b) Microglia activation in the DG and mPFC negatively correlated with prosocial behaviour

To measure the relationship between microglia activation and sociability, we ran a linear regression model including microglia activation, sex and OT/OTA/control with change in time spent in the social chamber during the sociability test from saline to LPS treatment as the outcome variable. Mice with greater microglial activation in the DG had reduced sociability following LPS relative to saline, regardless of OT/OTA/control group (b = −174.38 ± 73.49, p = .023, F(4, 38) = 2.66, R^2^ = .22, adj. R^2^ = .13) (Figure 3.A). There was no sex interaction effect on microglial activation in the DG and change in sociability (p = .635) (Supplemental Figure 15.A), and no relationship between change in sociability and microglial activity in the mPFC (p = .124) (Figure 3.B) or MeA (p = .249) (Figure 3.C).

**Figure 3.**
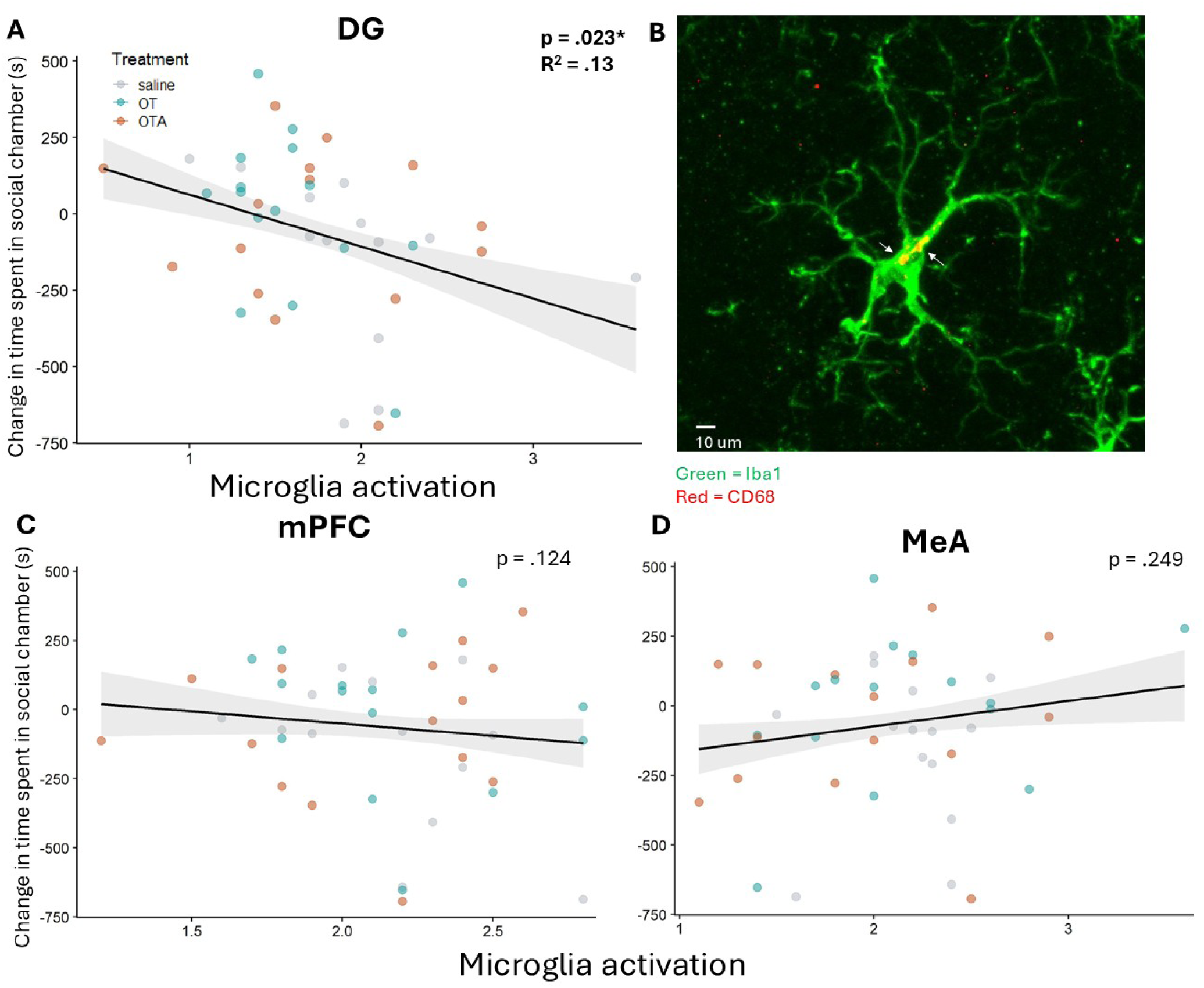
A) Mice with greater microglial activation in the dorsal DG spent less time in the social chamber following LPS relative to when they received saline, regardless of treatment condition (p = .032, R^2^ = 1.9, adj. R^2^ = .10). B) Representative image of a single microglia with two CD68+ inclusions indicated by white arrows at 20x magnification. There was no association between microglial activation in the mPFC (C) (p =.124) or MeA (D) and change in time spent in the social chamber (p = .249).

We also ran a linear regression model including microglia activation, sex and OT/OTA/control with time in proximity during undisturbed observations as the outcome variable while statistically controlling for baseline proximity. Mice that showed greater proximity either after saline (b = −60.05 ± 20.04, p = .010, F(5, 35) = 4.61, R^2^ = .40, adj. R^2^ = .31) or LPS (b = −78.79 ± 30.67, p = .014, R^2^ = .25, adj. R^2^ = .15) had reduced mPFC microglia activation at day 9. (Figure 4.B). There was no significant difference between the relationship between mPFC microglia activation and change in proximity following LPS relative to saline (p = .94). There was no sex difference in the relationship between microglial activation in the mPFC and time in proximity after saline (p = .091) (Supplemental Figure 15.B, C) or LPS treatment (p = .124).

**Figure 4.**
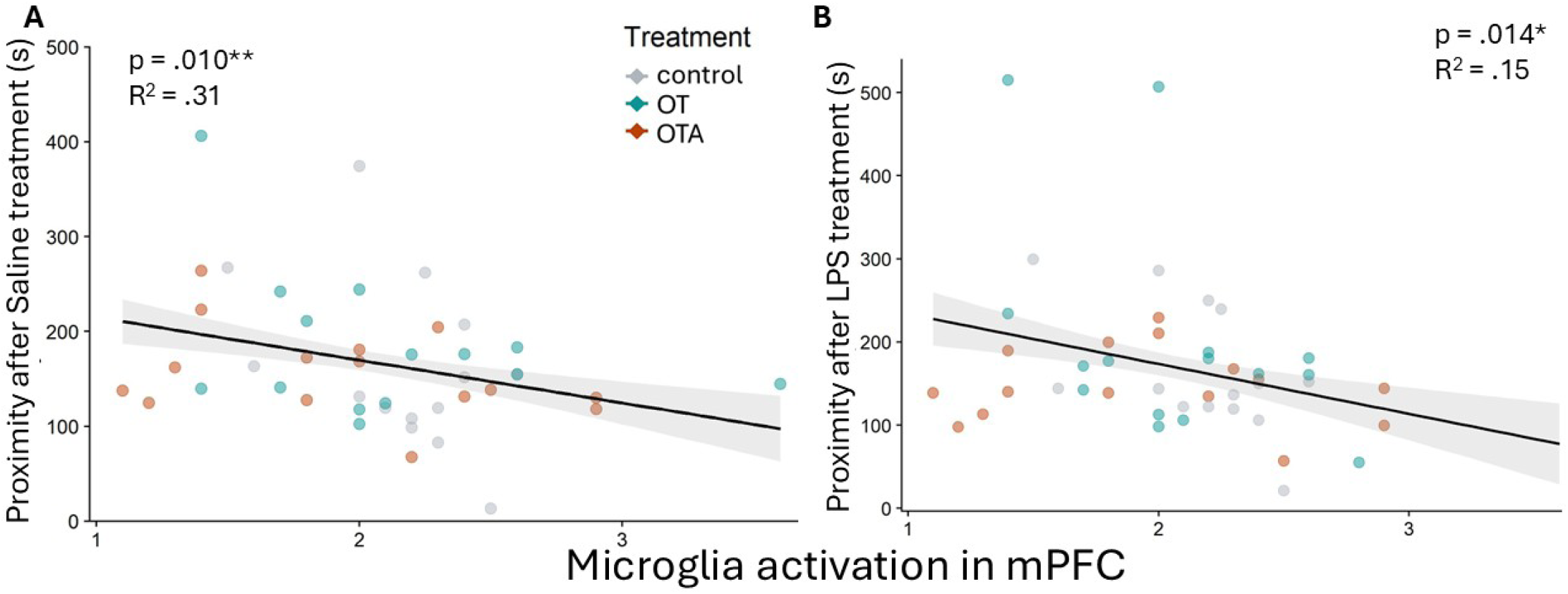
Mice with greater microglial activation in the mPFC spent less time in close proximity. Mice with greater microglial activation in the mPFC spent less time in close proximity after both A) saline (p = .010, R^2^ = .40, adj. R^2^ = .31) and B) LPS (Day 7) (p = .014, R^2^ = .25, adj. R^2^ = .15). P-values displayed are Holm-Bonferroni corrected for multiple comparisons.

### (c) Elevated proinflammatory cytokine release was associated with reduced microglia activity in the mPFC and time spent in close proximity

We ran a linear regression model and regressed PC1, PC2, PC3, and OT/OTA/control on microglial activation controlling for sex. PC1, associated with proinflammatory cytokines, negatively correlated with microglia activation in the mPFC (b = -.112 ± .041, p = .012, F(6,29) = 2.28, R^2^ = .32, adj. R^2^ = .18) (Figure 5.A) with no effect of OT (p = .512), OTA (p = .445), PC2 (p = .951) or PC3 (p = .230). PC1 did not correlate with microglial activation in either the DG (p =.713) (Supplemental Figure 11.A) or MeA (p = .720) (Supplemental Figure 11.B). There was also no difference between sexes on the relationship between PC1 and mPFC microglial activation (p = .180) (Supplemental Figure 15.D). We ran a linear regression model and regressed PC1 and behavioural phenotype on proximity controlling for sex and OT/OTA/control treatment. PC1 positively correlated with time spent in close proximity following LPS in huddler mice of both sexes, but had no relationship in non-huddlers (b = 47.88 ± 13.65, p = .002, F(9,27) = 8.67, R^2^ = .74, adj. R^2^ = .66) (Figure 5.B).

**Figure 5.**
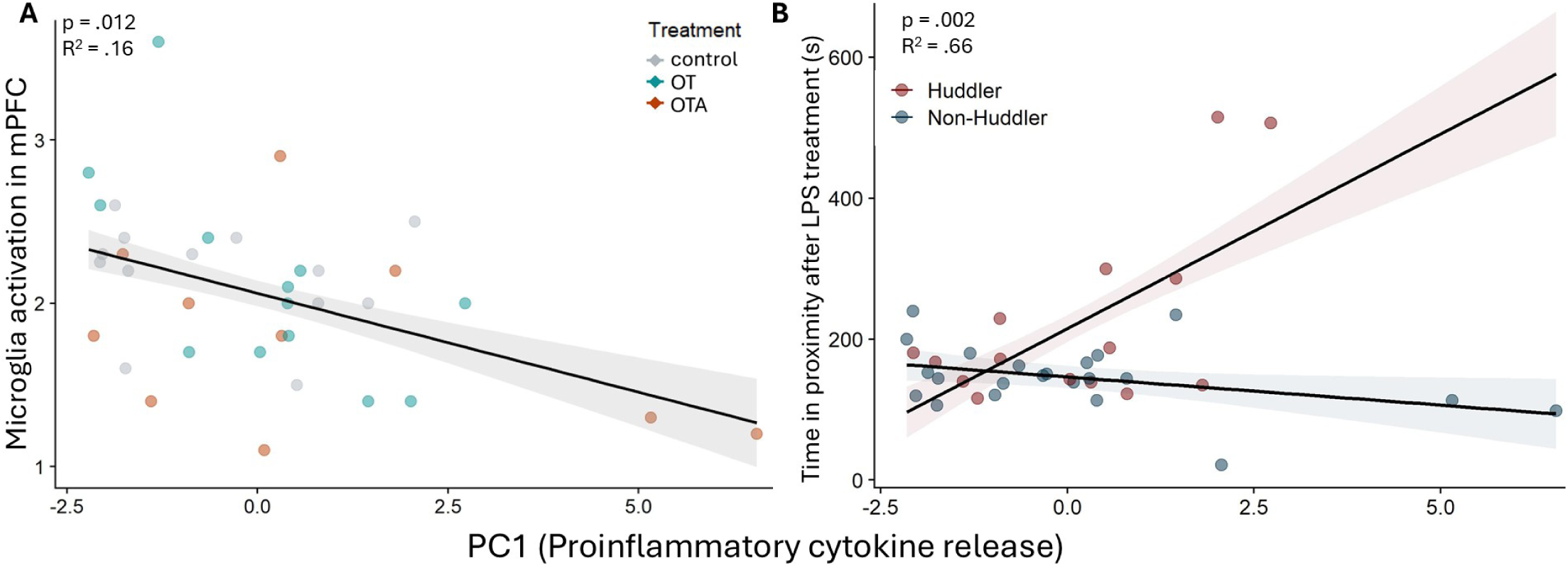
A) Peripheral cytokine release, represented by PC1, was negatively correlated with microglia activation in the mPFC (p = .012, R^2^ = .32, adj. R^2^ = .18). B) Female and male huddlers had a positive correlation between cytokine release and time spent in proximity following LPS (p = .002, R^2^ = .74, adj. R^2^ = .66).

### (d) LPS altered sweep call features and relationship between olfactory investigation and sweeps

Following LPS mice had longer sweep durations (b = .99 ± .26, p < .001, F(1,44) = 14.8, conditional R^2^ = .46, marginal R^2^ = .35), reduced power (b = −1.28 ± .47, p = .009, F(1,44) = 7.52, conditional R^2^ = .77, marginal R^2^ = .57), increased sinuosity (b = .36 ± .14, p = .015, F(1,41) = 6.39, conditional R^2^ = .24, marginal R^2^ = .16), and increased principle frequency (b = 5.10 ± 1.60, p = .003, F(1,41) = 10.13, marginal R^2^ = .45). Additionally, mice had sweep calls that were of a less steep negative slope after LPS (b = 602.66 ± 145.56, p < .001, F(1,41) = 17.41, conditional R^2^ = .49, marginal R^2^ = .47), and this effect was stronger in females (b = −600.96 ± 219.27, p = .009, F(1,41) = 7.51, conditional R^2^ = .49, marginal R^2^ = .47). Furthermore, we found that females had higher call frequencies compared to males regardless of treatment (b = 3.64 ± 1.61, p = .029, F(1,40) = 5.14, marginal R^2^ = .45). The only difference in behavioural phenotype observed was that control huddlers did not increase call frequency following LPS compared to OT/OTA mice or non-huddler controls (b = −9.45 ± 3.34, p = .008, F(1,41) = 7.89, conditional R^2^ = .59, marginal R^2^ = .52). Finally, we found that after LPS, but not saline, there was a positive relationship between olfactory investigation and call length (b = .011 ± .004, p = .013, F(1,70) = 6.5, marginal R^2^ = .36) and call frequency (b = 2.79 ± 1.00, p = .008, F(1,68) = 7.5, marginal R^2^ = .49). For details on LPS effects on vocalizations, see Supplemental Results 9.g, h.

## 4. Discussion

During an immune challenge, social animals must choose between affiliation and avoidance. While high doses of LPS induce sickness behaviours, social response to low-grade inflammation remains poorly understood. This is critical to understand because mild inflammation occurs not only during minor illness or injury but also in anxiety and depression [38,56]. We tested whether social-behavioural phenotype predicted social response to low-dose LPS, whether microglia activity mediates peripheral inflammation-induced changes in social behaviour, and whether OT modulates this relationship. We found that behavioural phenotype predicted affiliative response to LPS in males, and that microglia activation in the mPFC and DG correlated with affiliation and sociability in a region-specific manner.

### (a) LPS differentially affected huddler and non-huddler males

Behavioural phenotype predicted male behavioural response to LPS: non-huddlers increased sociability, whereas huddlers did not (Figure 2). Huddling promotes bonding and neural plasticity [57], and individual variation in affiliation is common across social species [41]. Our findings show this variation is key to predicting social decision-making during an immune challenge.

We predicted LPS would increase sociability in California mice due to their high affiliative capacity [10,16], and because same-sex proximity to a familiar individual improves healing [14]. In non-huddlers, LPS increased sociability (Figure 2), consistent with rat and rhesus macaque studies showing inflammation-induced affiliation [3,4]. We expected that more affiliative huddlers would have a greater increase in sociability following LPS, but we found no effect of LPS on sociability in huddlers. Huddlers may already be at a behavioural ceiling, or their motivation to affiliate is consistently high, while non-huddlers flexibly adjust behavior according to their immune state.

Interestingly, we saw no change in sociability following LPS in females across phenotypes. Although sex steroid hormones shape immune function [58], we found no sex differences in microglia activation or peripheral cytokines (Supplemental Figure 15). We therefore suggest females may be more behaviourally resilient to immune challenge rather than immunologically different. Similar sex-specific interactions between sociality and immunity have been reported, with group housing reducing inflammation in male but increasing it in female rats [4]. Together, these findings support the intriguing possibility that there is a positive relationship between sociality and immune function in males but not females, a hypothesis that warrants further exploration.

We predicted OT would enhance, and OTA reduce, sociability during inflammation based on OT’s prosocial and anti-inflammatory effects [21]. However, neither OT nor OTA influenced social decision making or immune outcomes, suggesting OT is not the primary mechanism linking immune function and social decision making. OT effects on social behaviour are known to be social context-dependent, demonstrated in California mice, rats, and humans [18,35,59], and often emerge under social defeat or isolation stress [14,35], conditions absent in this study. OT remains an important candidate mechanism for mediating social-immune interactions, but was not the driver of behaviour changes observed in this study. Instead, we propose that elevated proinflammatory cytokines during immune activation or stress may suppress microglial activity in the mPFC, and mPFC microglia regulate social responses to immune change (Figure 6).

**Figure 6.**
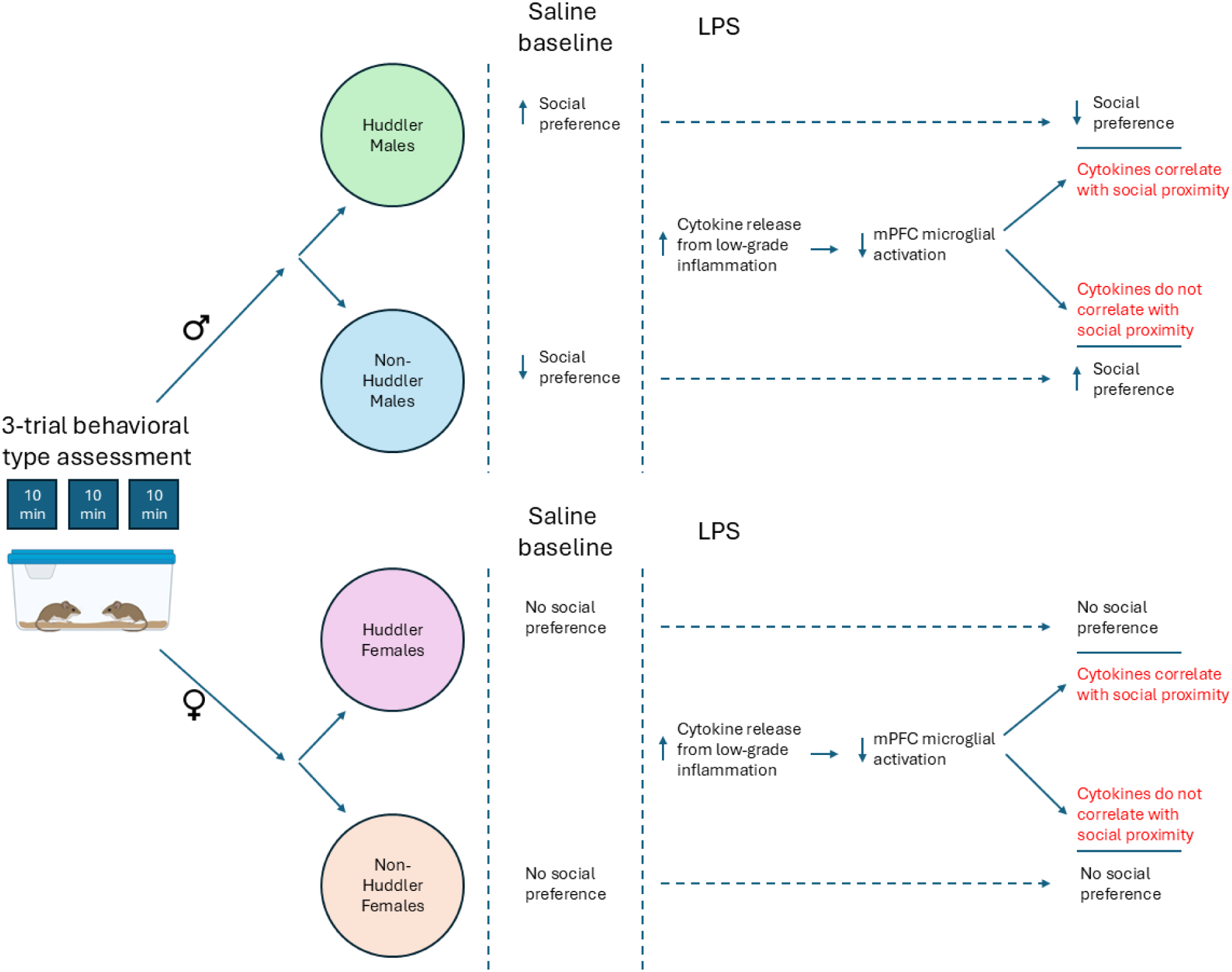
Proposed mechanism of action and results summary. We show two main findings: 1) Huddler versus non-huddler behavioural phenotype predicted social response to LPS. 2) We revealed correlations between peripheral cytokines, mPFC microglia activation, and social proximity that suggest that proinflammatory cytokines may increase affiliation through inhibition of mPFC microglia, and an important future direction is to test this experimentally.

### (b) Microglia and cytokines may regulate social decision-making

Microglia shape synaptic pruning and plasticity, making them integral to social behaviour [24,25]. We predicted that increased microglial activation would be associated with reduced affiliation during immune challenge. Consistent with this prediction, higher microglial activation in the mPFC was associated with reduced proximity to a familiar mouse, and activation in the DG was associated with reduced sociability; no association was found with the MeA. This aligns with evidence that increased microglial activation in the mPFC and DG decreases sociability and increases depressive-like behaviour via excessive pruning and altered synaptogenesis [60,61]. Our results are correlational, and the negative relationship between mPFC activation and proximity was present after both saline treatment and LPS, suggesting this relationship is present even in non-inflamed conditions (Figure 4). Another possibility is that greater proximity leads to reduced microglia activity in the mPFC. This would be consistent with evidence that prosocial behaviour may be immunoprotective [62]. Regardless, our results highlight the mPFC as a key region where microglia influence social behaviour.

Because proinflammatory cytokine activity is typically associated with elevated microglia activity [63], we expected that DG and mPFC activation would positively correlate with microglia activation. Intriguingly, we found mPFC microglial activity negatively correlated with peripheral cytokines, with no relationship in the DG or MeA (Figure 5.A; Supplemental Figure 11). This supports region-specific microglial responses [64]. These regional differences may have behavioural implications; while stress broadly increases microglial activation [65] both non-social stress [66,67] and social stress [68] reduce microglial activation in the mPFC specifically. As stress is associated with elevated peripheral cytokine release [69], we speculate that many cytokines considered “proinflammatory” may inhibit mPFC microglia during physiological or social stress, though this needs to be experimentally tested. We propose that cytokines can inhibit mPFC microglial activation, thereby disinhibiting affiliation (Figure 6).

Behavioural phenotype also shaped the relationship between the immune system and social behaviour: in huddlers, higher proinflammatory cytokine expression correlated with greater proximity, whereas non-huddlers showed no such relationship (Figure 5.D). This suggests that inflammation may enhance affiliation through inhibition of mPFC microglia, depending on behavioural phenotype. The mPFC is well-positioned to mediate the relationship between immune state, social relationships, and behavioural decision-making, as neural circuitry in the mPFC is necessary for social perception, social learning, and a context-appropriate social response [31].

Because of their ability to cross the blood-brain-barrier, it was proposed that cytokines serve an important social function for modulating social behaviour in both pathological and healthy conditions [70]. However, there may be differences between peripheral and central cytokine levels. As we measured cytokines peripherally, we cannot rule out that the cytokine profile in the mPFC was different from the plasma. As the primary immune cells in the brain, microglia may be the critical link between cytokine activity and social function. As our data are correlational, the relationship between proinflammatory cytokine activity and microglia activity in the mPFC is an important area for further exploration.

We hypothesize that microglia in the mPFC contribute to social decision-making, and are particularly important during an immune challenge. The absence of relationships in the MeA suggests region specificity. Causal tests will require region-specific microglial manipulation. In the DG, microglial activation negatively correlated with sociability and showed a non-significant trend toward OT reducing activation (Supplemental Figure 9), consistent with reports that DG microglial overactivation drives depressive-like avoidance behaviour and blocking this reverses social withdrawal [71].

Our goal was to evaluate individual variation in sociability under non-inflammed versus low-grade inflammatory conditions induced by LPS. However, because all mice received LPS prior to tissue collection, baseline microglial activity could not be assessed. Additionally, tissues were collected four days after LPS rather than at peak inflammatory activity (24 hrs). Nonetheless, the observed correlations between microglia, cytokines, and social behaviour provide insight into how central and peripheral immune systems interact to shape social function.

### (c) LPS altered olfactory social investigation and vocalizations

We explored whether LPS affected vocalizations and olfactory investigation. LPS altered multiple acoustic features of sweep calls: duration, slope, frequency, sinuosity, and power (Supplemental Figure 12). Prior studies show that LPS increases 22 kHz distress vocalizations in rats [73] and reduces social contact calls in vampire bats [73].To our knowledge, this is the first study to assess how LPS influences detailed acoustic features beyond call number alone. The descending “sweep” call, a short high-frequency vocalization associated with affiliation in California mice [15,16], increased in both average frequency and duration following LPS (Supplemental Figure 12). Although little is known about the behavioural implications of changes in distinct vocal features of sweep vocalizations, higher frequency and longer duration sweeps are associated with increased social contact in the closely related *Peromyscus truei* [74], suggesting LPS may promote more affiliative vocal signals. Additionally, LPS + OT reduced call power (i.e., amplitude) relative to control; lower amplitude calls are associated with closer social contact [75] These changes allow mice to communicate their “sick” or inflamed state, with longer sweeps being more affiliative.

These changes in vocalizations following LPS correlated with behavioural changes. LPS led to a positive correlation between olfactory investigation by the focal mouse and sweep duration (Figure 7). Olfactory investigation by mildly sick animals may reflect increased social approach or perhaps assessment of partner responses. Behavioural phenotype did not predict LPS-induced changes in most call features (Results 3.d), suggesting that while behavioural phenotype predicts social change following LPS in some measures, it does not determine how LPS influences vocal communication.

## 5. Conclusions

This study shows that 1) behavioural phenotype shaped immune-related sociability in males, with non-huddler males increasing sociability and no effect in huddler males; 2) microglial activation in the mPFC and DG was associated with reduced social behaviour, 3) OT did not mediate immune-induced behavioural changes, and 4) LPS altered vocal features, including increasing sweep duration and frequency, and longer sweeps were associated with increased olfactory investigation. Together, these findings deepen our understanding of the interactions between immune function and social behaviour, and provide a novel perspective on how behavioural type and prior affiliation may influence response to an immune challenge, as well as the glial mechanisms involved in this relationship.

## 6. Acknowledgements

Research was conducted at the University of Wisconsin-Madison, which occupies the ancestral Ho-Chunk land known as Teejop. Federal and state governments repeatedly, but unsuccessfully, sought to forcibly remove the Ho-Chunk from Wisconsin. As members of a land grant institution we directly benefit from land theft, and we challenge ourselves and others to reflect on the perpetuation of the colonialist roots of western scientific progress. Microscopy was performed at the Newcomb Imaging Center, Department of Botany, University of Wisconsin – Madison. We thank L. Riters for discussions on data analysis that strengthened this manuscript. We thank X. Zhou, A. Scapple, and N. Cangelosi for assisting in data collection. We also thank the UW-Madison animal research technicians. Research was supported by HHMI (GT16916), NSF (IOS-1946613 and DGE-1747503), and the Wisconsin Alumni Research Foundation at UW-Madison.

## 8. Supplemental Methods

### (a) Behaviours for undisturbed observations

**Supplemental Table 1.** Behavioural ethogram.

| Behaviour Name | Behaviour Description |
| --- | --- |
| Sniff given | The animal touches its nose any body part of its cagemate |
| Sniff received | The animal is touched by the nose of its cagemate on any body part |
| Huddling | The flanks of the two mice are in constant contact for greater than |
|  | 5 seconds |
| Proximity | Animals are within one mouse-length of each other for greater than 5 seconds |
| Flipping | The animal flips in its cage |
| Freezing | The animal is completely still and alert, ears up |
| Inactivity | The animal is not moving and not alert, ears are not turned upwards |

### (b) Huddling Analysis

Chi-squared analyses revealed no difference in the number of huddlers or non-huddlers across groups (p = .282) or sex (p = 1.0). A Chi-squared analysis was performed on huddling versus non-huddling at each time point: at baseline (X^2^ = 1.8, p = .18), after saline (X^2^ = 0, p = .87), and after LPS (X^2^ = 2.2, p = .14) (Supplemental Figure 1), demonstrating that the population maintained a consistent proportion of huddlers and non-huddlers. We therefore treat the huddler and non-huddler behavioural categories as consistent populations. Statistics evaluating huddling as a continuous variable were run on huddlers only (n = 18) after a log2 transformation to meet model assumptions; there was no significant effect of OT (p = .88), OTA (p = .38), LPS (p = .85), or sex (p = .25) were found on huddling behaviour.

**Supplemental Figure 1.**
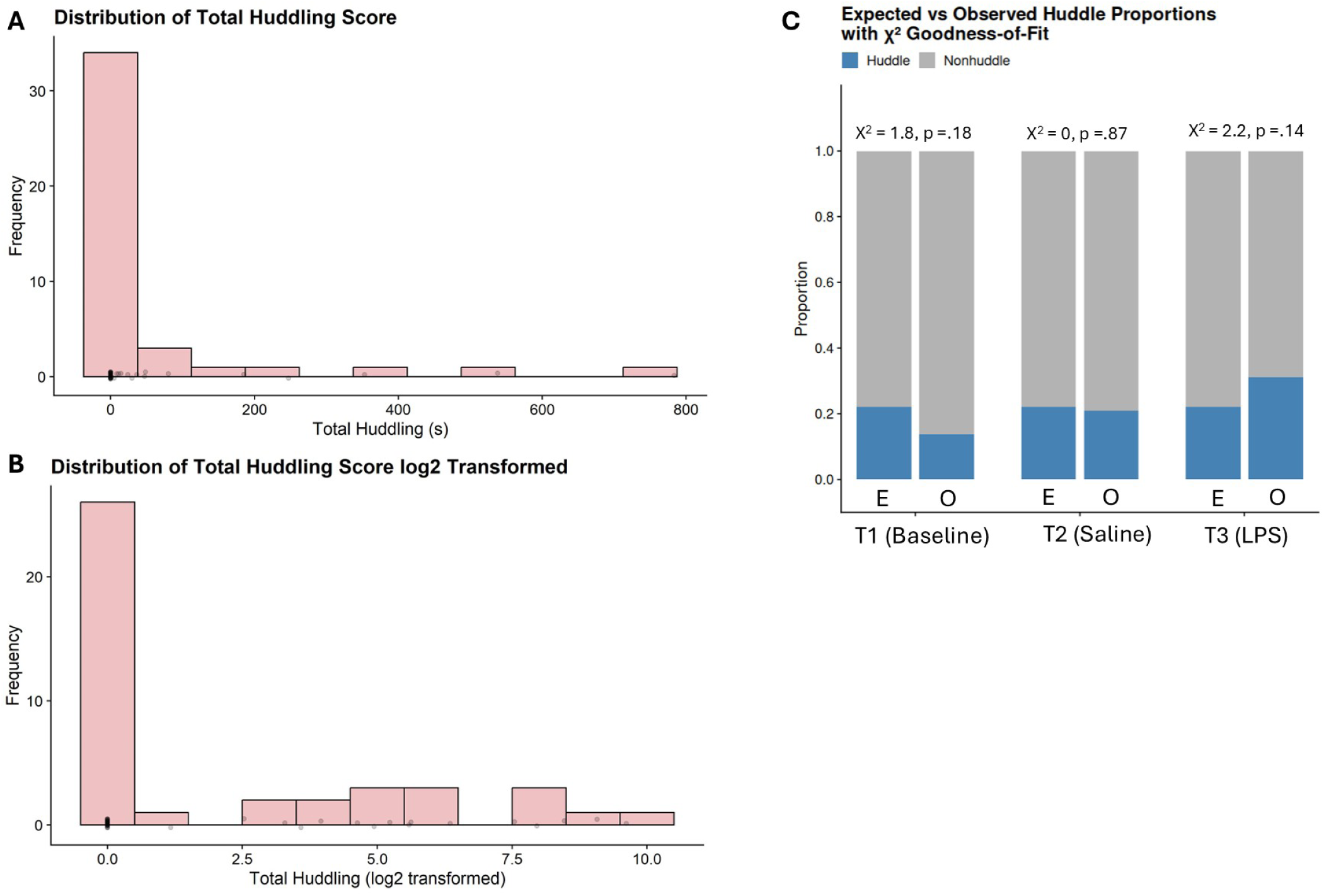
A) A histogram of raw huddling scores collected across all three 10-min observation periods. B) A histogram showing the distribution of total huddling score (log2 transformed) across all three 10-min observation periods. C) A chi-squared analysis showing expected (E) versus observed (O) behavioural phenotype. At T1 (baseline) X^2^ = 1.8, p = .18. At T2 (saline) X^2^ = 0, p = .87. At T3 (LPS) X^2^ = 2.2, p = .14.

### (c) Elevated Plus Maze Test

During the elevated plus maze (EPM) test, the mouse was placed in a four-armed EPM raised 1 meter off of the ground. Two arms were closed, and two arms were open, each arm 67-cm long and 5.5-cm wide. Tests lasted 5-min, and time spent in open arms and entries into closed arms were recorded. No changes were observed across treatment during the EPM (p = .351), suggesting that the effects demonstrated are not a result of differences in anxiety-like behaviour (Supplemental Figure 6).

### (d) Microglia Immunocytochemistry

The day following the elevated plus maze test and third treatment of OT/OTA/control, mice were deeply anesthetized with isoflurane and transcardially perfused with 4 % PFA in PBS. Brains were fixed for 48 h in 4% PFA followed by cryoprotection with 30% sucrose in PBS for 48 h. Unilateral 40 μm-thick coronal sections embedded in OCT were cut using a cryostat (Leica Biosystems). Sections were washed with PBS, blocked with 5% normal donkey serum, and permeabilized with 0.5 % Tween-20 in PBS for 1 h at room temperature. Polyclonal rabbit anti-Iba1 (Avantor catalog #100369-764) and monoclonal rat anti-CD68 (Abcam catalog #ab53444) primary antibodies were added for 24 hrs at room temperature at a dilution of 1:500 and 1:200, respectively. After washing, sections were incubated with donkey anti-rabbit AlexaFluor-488 (Abcam catalog #ab150073) and donkey anti-rat AlexaFluor-568 (Abcam catalog #ab175475) conjugated secondary antibodies for 1.5 h in the dark at dilutions of 1:250. Sections were rinsed, mounted onto Suprafrost Plus slides (Thermo Fisher Scientific), dried, counterstained with Hoechst 33342 (Invitrogen) 1:1000 in water, and coverslipped using Immu-Mount (Thermo Fisher Scientific).

**Supplemental Figure 2.**
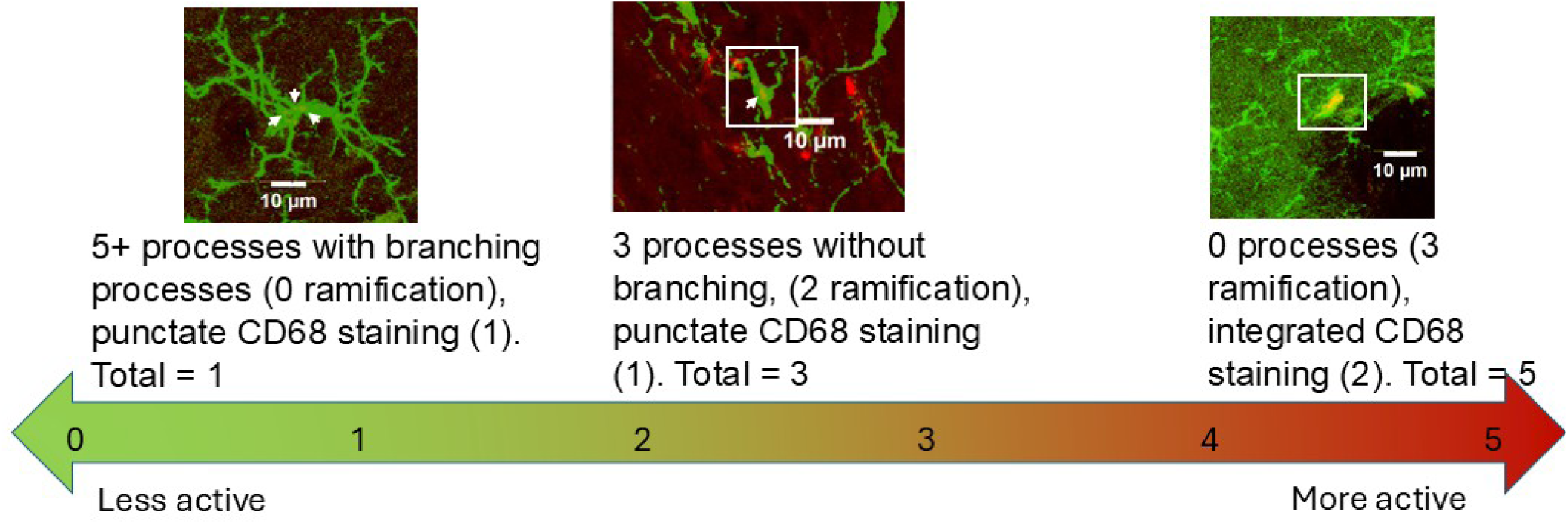
Microglia activation state was categorized on a 0 (lowest activation) to 5 (highest activation). Morphology, identified through Iba1 staining, was scored as 0 (5+ processes with at least secondary branches), 1 (1 - 4 processes with at least secondary branches), 2 (1+ processes with no secondary branches), and 3 (round with no clear processes). CD68 expression was scored as 0 (no expression), 1 (punctate expression) or 2 (aggregated expression). Morphology and CD68 scores were then combined to get a final score from 0-5.

**Supplemental Figure 3.**
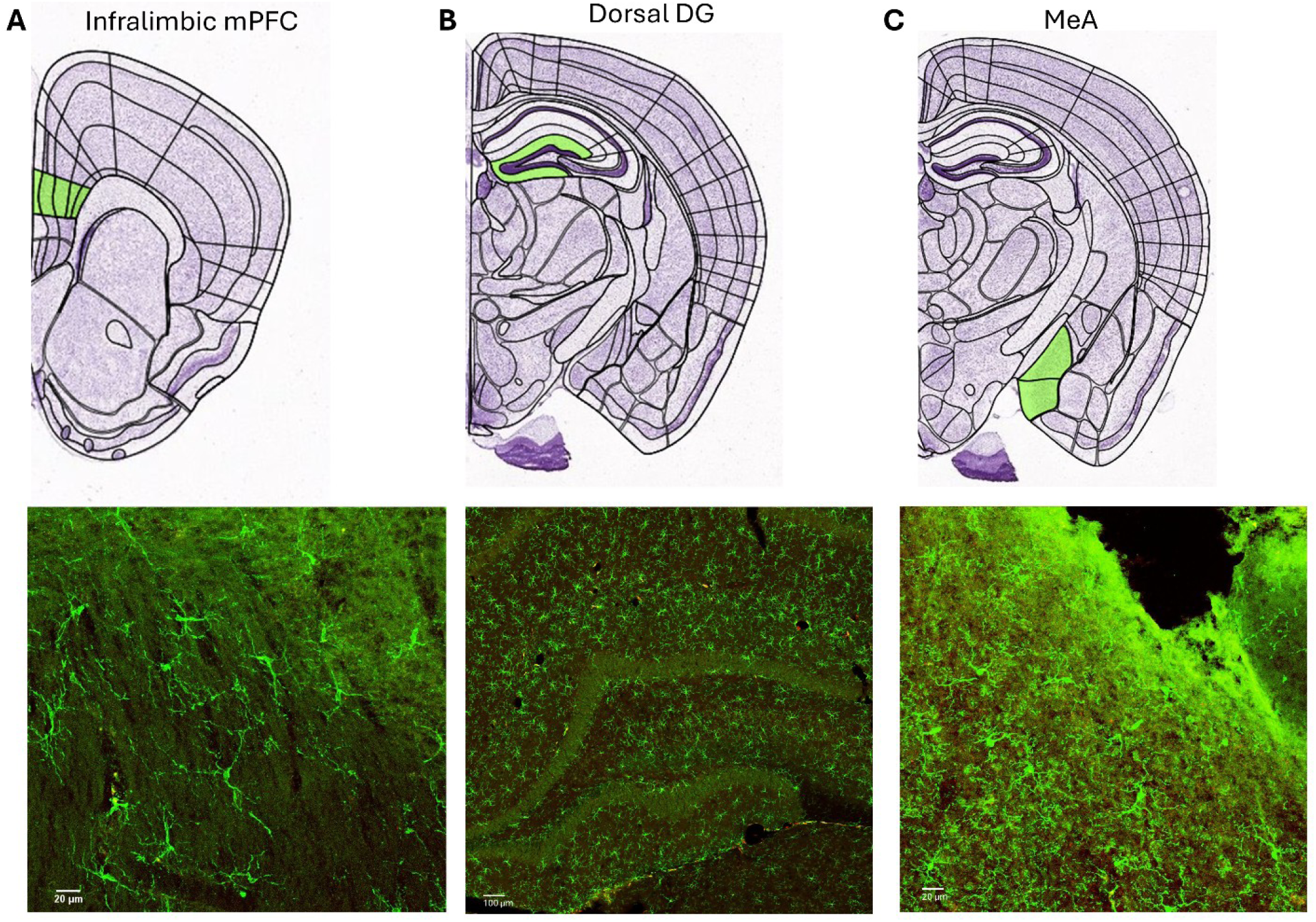
Graphical depiction of where confocal images were taken from the brain. Images used from the Allen Brain Atlas, as well as representative confocal images of microglia staining in these brain regions.. A) Green highlights the infralimbic region of the mPFC. Image taken at 20x magnification. B) Green highlights the dorsal region of the DG. Image taken at 10x magnification. C) Green highlights the MeA.

### (e) Electrochemiluminescence

As proinflammatory Panel 1 (mouse) plate comes precoated with the analyte’s capture antibodies, the plate was briefly washed with a Agilent BioTek 405TS Microplate Washer three times with a manufactured MSD wash buffer. Samples stored at −80°C were thawed and diluted 1:2 with MSD dilution buffer. 50uL of sample was added to each well in duplicates and incubated for 2-h at room temperature with shaking. After incubation, the plate was washed three times with MSD buffer to remove any unbound antigens. Following this, a solution of each analyte’s detection antibody, conjugated with “SULFO-TAG” electrochemiluminescent labels, was added to the plate and incubated for 2-h at room temperature with shaking. The plate was then rinsed three times with MSD buffer. Finally, 50ul of Read Buffer T was added to each well and immediately quantified with the MESO QuickPlex SQ 120. This instrument uses electrodes to interact with the electrochemiluminescent labels of the SULFO-TAG antibodies to measure the concentration of the 10 analytes within each well based on the intensity of signal emitted according to the standard curves of each analyte. Standard curves used for calculating concentration had R^2^ values of .99.

**Supplemental Figure 4.**
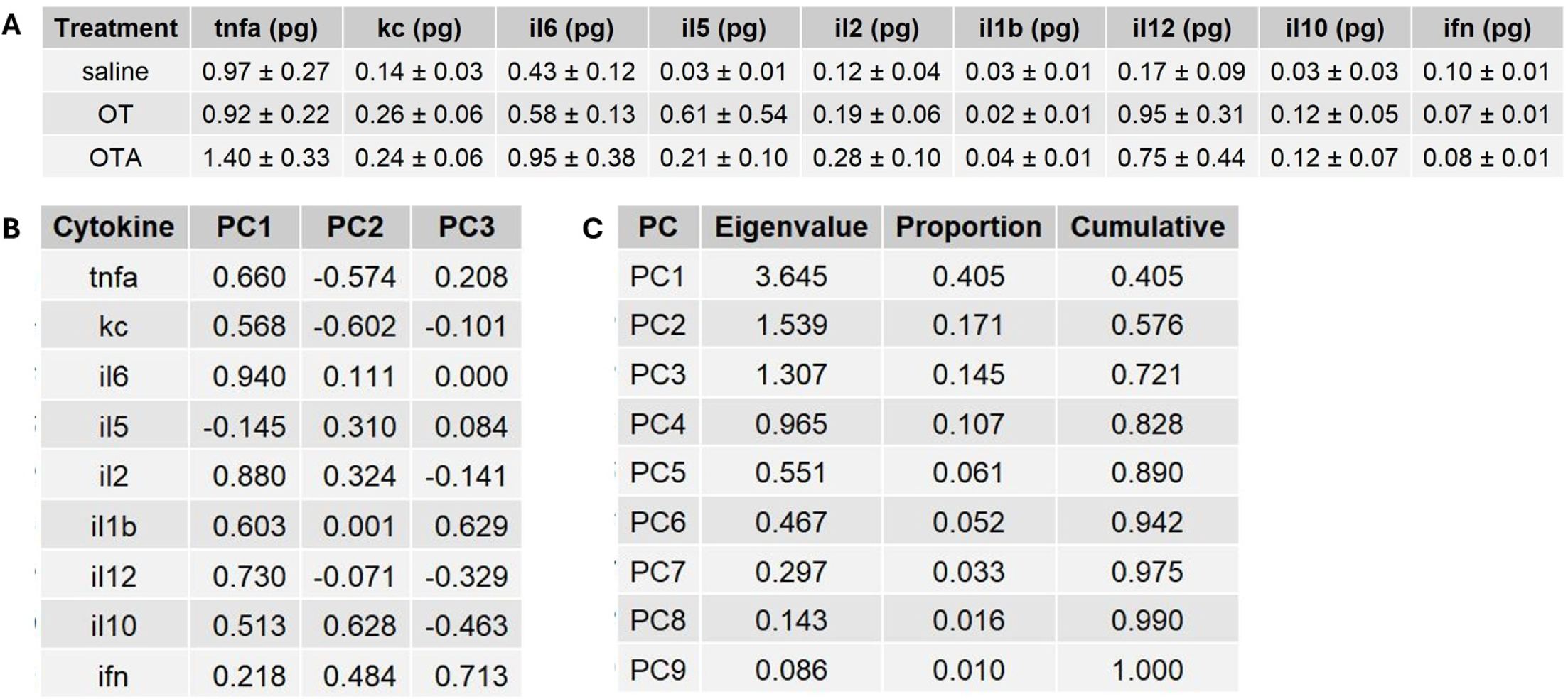
A) The means and SEMs of cytokines detected via electrochemiluminescence by treatment group. There were no significant differences across groups. B) Top three principal component correlation loadings for the nine measured cytokines following a principal component analysis. Cytokines measured included tumor necrosis factor alpha (TNF-ɑ), keratinocyte chemokine (KC), interleukin-6 (IL-6), interleukin-5 (IL-5), interleukin-2 (IL-2), interleukin-1 beta (IL-1b), interleukin-12 (IL-12), interleukin-10 (IL-10) and interferon gamma (IFN). CB) The Eigenvalues, proportion of variance explained, and cumulative variance explained by each principal component.

**Supplemental Figure 5.**
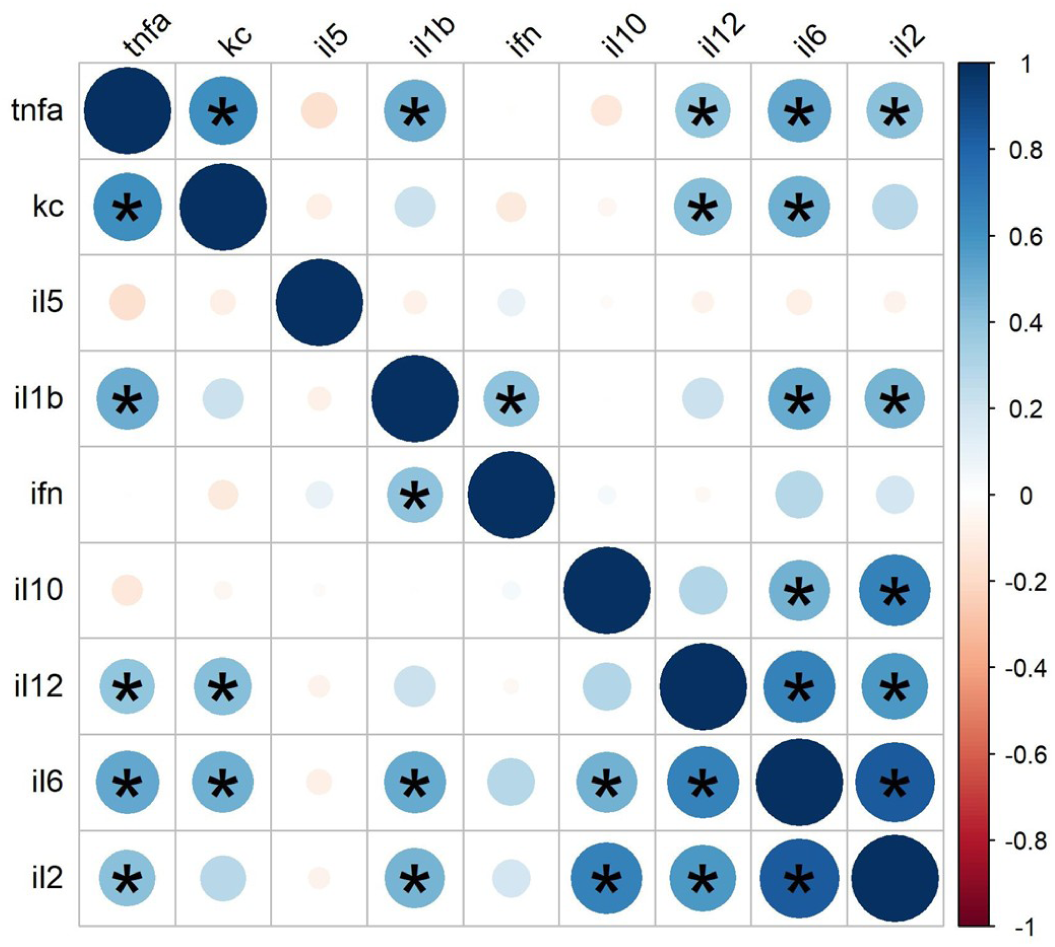
Correlation matrix of raw values of cytokines included in this study. An asterisk indicates a correlation of p < .05.

## 9. Supplemental Results

### (a) Changes in activity and weight following LPS

We ran a mixed effects regression model including OT/OTA/control treatment the between-subject variable, and LPS/saline as the within-subject variable, and time spent inactive as the outcome variable. Mice did not spend more time inactive as a result of LPS in any treatment group (p = .504).

**Supplemental Figure 6.**
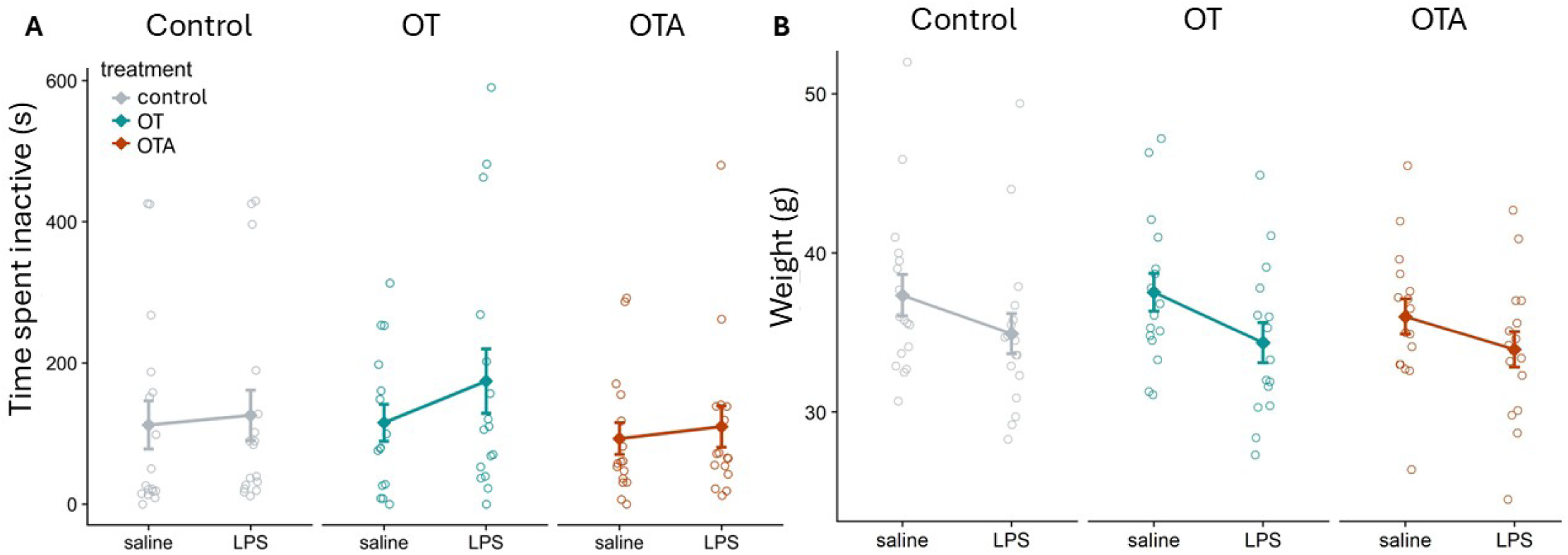
A) Mice did not spend more time inactive as a result of LPS in any treatment group (p = .504). B) Mice weighed significantly less following LPS, losing an average of 2.40g ± 4.06 throughout the duration of the study (p < .001, F(1,42) = 34.96, conditional R^2^ = .94, marginal R^2^ = .07). There was no difference between control and OT (p = .97) or control and OTA (p = .50).

### (b) EPM Results

We ran mixed effects regression models including sex and OT/OTA/control treatment the between-subject variables, and LPS/saline as the within-subject variable, and time spent in the open arm and number of passes into the closed arm as the outcome variables.

**Supplemental Figure 7.**
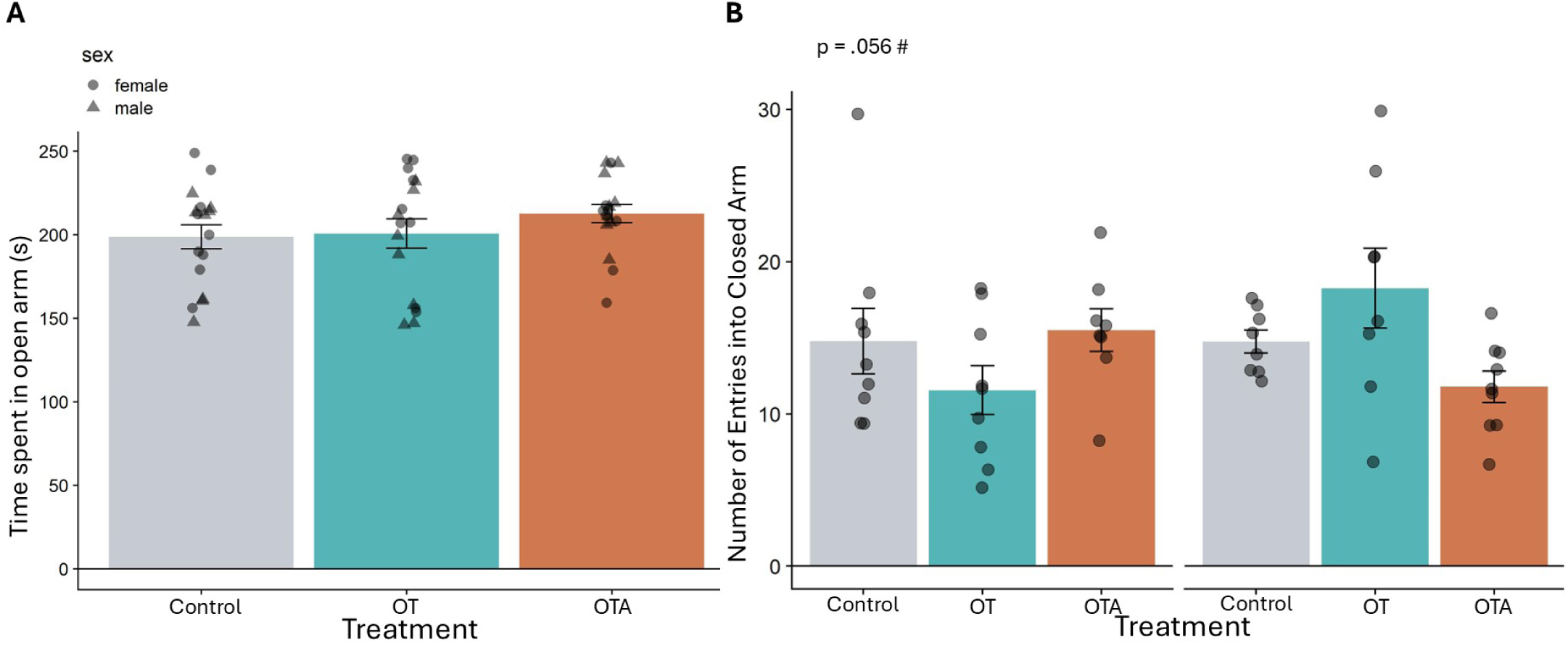
A) There were no differences in time spent in the open arm of an elevated plus maze as a result of treatment (p = .351). B) There was a trending interaction effect of OT and sex on number of entries into the closed arm (b = −6.72 ± 3.43, p = .056, F(5,45) = 2.13, R^2^ = .19, adj. R^2^ = .08), with no simple effect of OT on males (p = .168) or females (p = .178).

### (c) Oxytocin and oxytocin antagonist treatment reduced cagemate social investigation following LPS

We ran a linear regression model and regressed sex and OT/OTA/control on change in sniffing behaviour of the focal mouse and cagemate following saline and following LPS, and used Holm-Bonferroni corrections for multiple comparisons. OT (p = .33) and OTA (p= .23) alone did not influence the number of times a mouse was sniffed by its cagemate (Supplemental Figure 7.A). Interestingly, following LPS, mice that received either OT or OTA were more frequently sniffed by their same-sex cagemate (Supplemental Figure 7.B). OT decreased sniffing done by the cagemate compared to control by an average of 3.90 sniffs ± 1.57 (p = .017, F(5, 40) = 2.05, R^2^ = .20, adj. R^2^ = .10). OTA also decreased sniffing by an average of 4.10 sniffs ± 1.47 (p = .008, F(5, 40) = 2.05, R^2^ = .20, adj. R^2^ = .10) relative to control. There was no effect of sex (p = .44) or an interaction between sex and treatment (p = .30 for OT:sex interaction, p = .68 for sex:OTA interaction). There was no change in sniffing behaviour of the focal mouse in control (p = .878), OT (p = .752), or OTA-treated mice (p = .944), demonstrating that the behavioural changes observed in a dyadic in which one animal is sick may often be driven by changes in behaviour of the healthy animal.

**Supplemental Figure 8.**
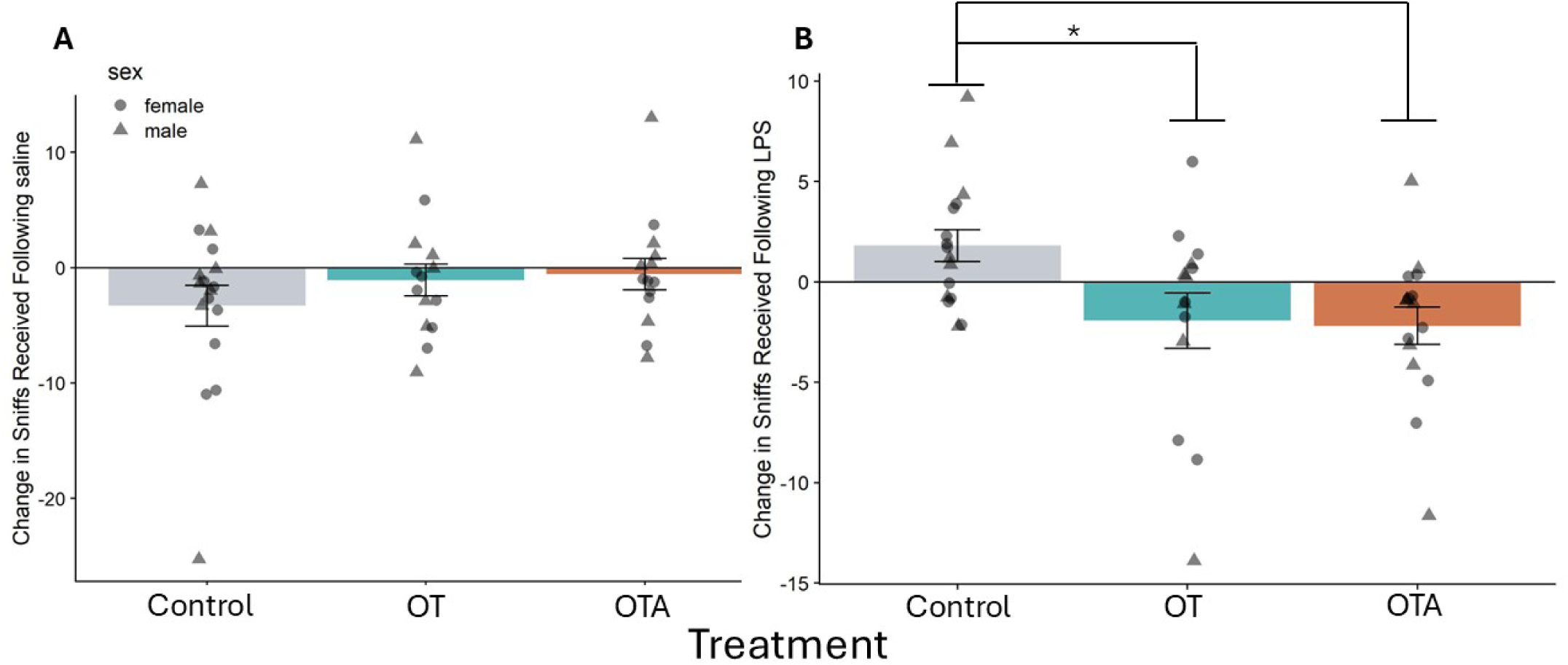
A) OT (p = .33) and OTA (p= .23) alone did not influence the number of times an animal was sniffed by its cagemate. B) OT treated mice were sniffed less following LPS, (p = .017, R^2^ = .20, adj. R^2^ = .10) as were OTA treated mice (p = .008, R^2^ = .20, adj. R^2^ = .10).

The similarity in response to OT versus OTA are likely driven by different mechanisms. It is possible that treatment with OT enhances communication of a distressed or sick state, leading to increased avoidance from the partner, while OTA may have disrupted the social behaviour of the focal mouse, similarly leading to cagemate avoidance.

### (d) OT trended towards reducing microglia activity in the DG, and did not alter microglia activation in the medial amygdala or mPFC

We ran a linear regression model and regressed sex and OT/OTA/control on microglia activation. Mice treated with OT had a nonsignificant trend towards reduced microglial activation in the dorsal DG relative to controls (b = -.040 ± .204, p = .059, F(3, 39) = 1.39, R^2^ = .10, adj. R^2^ = .03). There was no effect of OTA in the DG (p = .218). There was no effect of OT in the MeA (p = .912) or mPFC (p = .907), and no effect of OTA in the MeA (p = .659) or mPFC (p = .264) (Supplemental Figure 9).

**Supplemental Figure 9.**
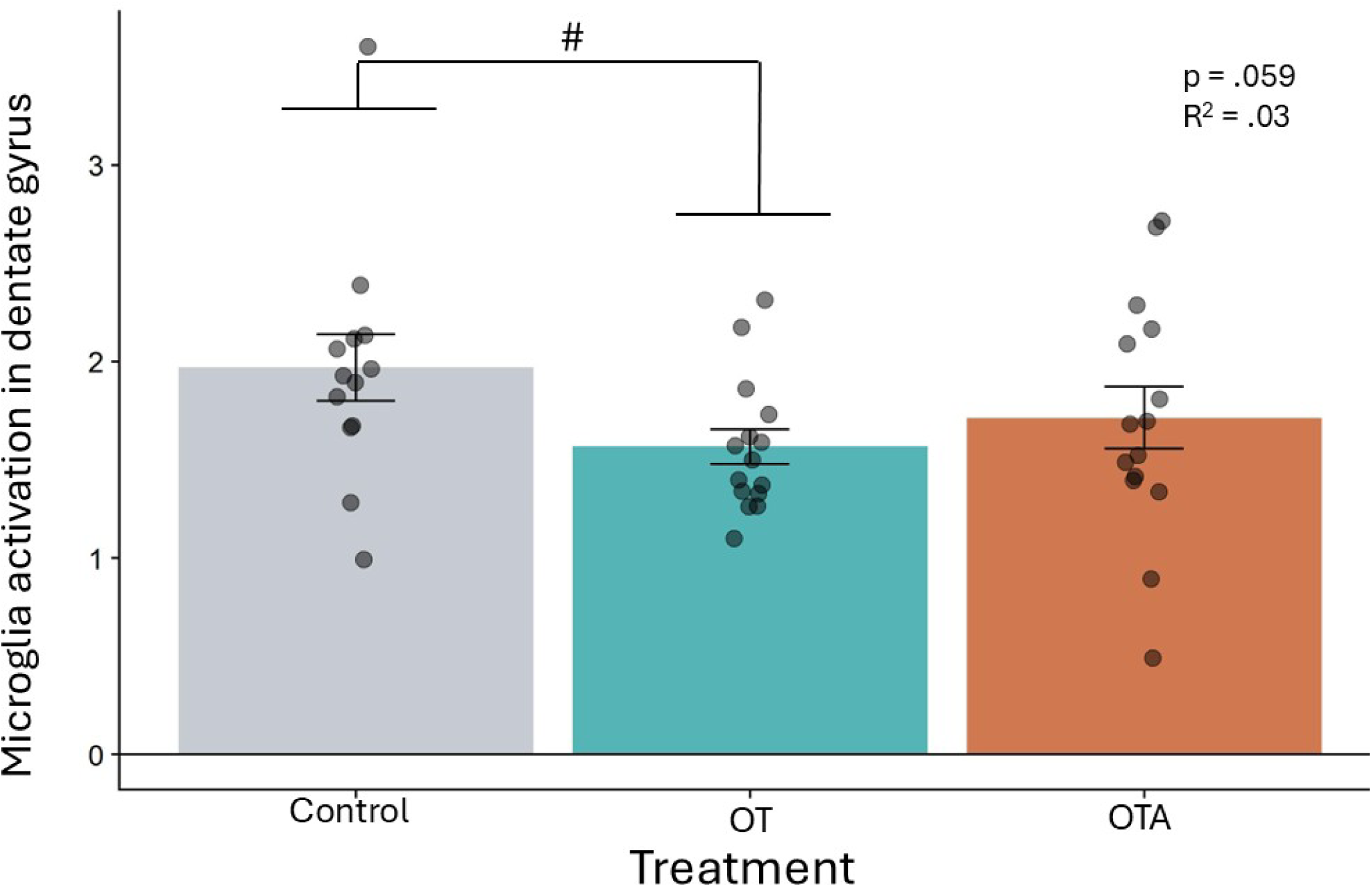
Mice treated with OT trended towards having reduced microglial activation in the dorsal DG relative to control (p = .059, R^2^ = .10, adj. R^2^ = .03).

### (e) Behaviour correlations

**Supplemental Figure 10.**
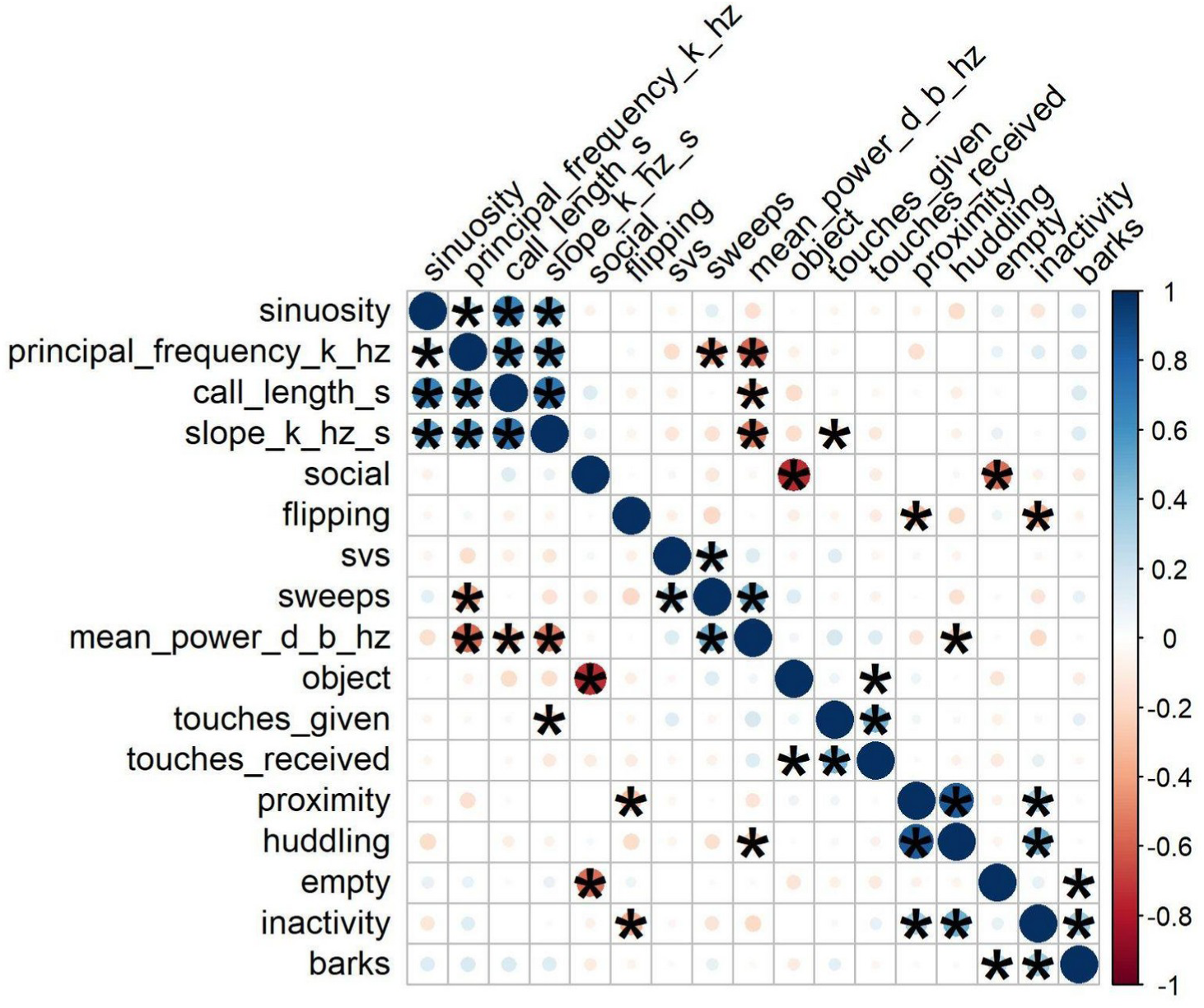
A correlation matrix showing all behaviours measured. An asterisk indicates a significance of p < .05.

### (f) There was no effect of cytokine release and microglia activation in the DG or MeA

**Supplemental Figure 11.**
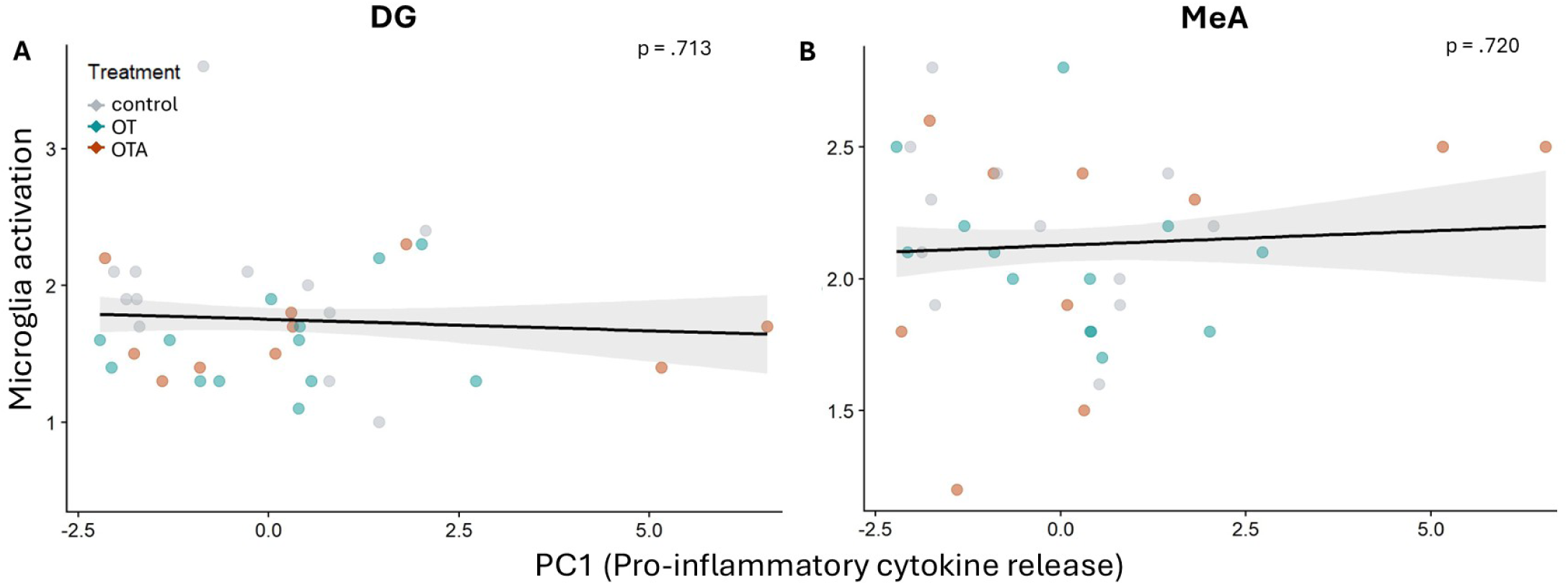
A) There was no relationship between DG (p = .713) or B) MeA microglial activation (p = .720) and PC1.

### (g) Sweep vocalizations increased in frequency and length, and reduced power following LPS

We ran a mixed effects regression model including behavioural phenotype, sex, OT/OTA/control treatment as between-subject variables and baseline vocalization data from day 1 as statistical covariates, and LPS/saline as a within-subject variable and vocal features categorized by DeepSqueak as the outcome variables. All vocal features of sweep calls were averaged for each individual cage. Following LPS, sweep call length increased relative to saline (b = .99 ± .26, p < .001, F(1,44) = 14.8, conditional R^2^ = .46, marginal R^2^ = .35). (Supplemental Figure 12.A). There was no effect of behavioural phenotype (p = .68) or sex on call length (p = .35) (Supplemental Figure 15.E). Mice vocalized with reduced relative power (dB/Hz) following both LPS (b = −1.28 ± .47, p = .009, F(1,44) = 7.52, conditional R^2^ = .77, marginal R^2^ = .57) and OT (b = −1.24 ± .56, p = .033, F(1,44) = 4.85, conditional R^2^ = .77, marginal R^2^ = .57) when compared to saline-treated mice (Supplemental Figure 12.B). There was no effect of behavioural phenotype (p = .32) or sex on average call power (p = .56) (Supplemental Figure 15.F). The average slope (kHz/s) of descending sweep calls became less negative, or less steep, following LPS (b = 602.66 ± 145.56, p < .001, F(1,41) = 17.41, conditional R^2^ = .49, marginal R^2^ = .47) (Supplemental Figure 12.C). Additionally, there was an interaction effect between OT and sex, such that females treated with OT had more negatively sloped calls (b = −600.96 ± 219.27, p = .009, F(1,41) = 7.51, conditional R^2^ = .49, marginal R^2^ = .47) (Supplemental Figure 12.C). There was no effect of behavioural phenotype on call slope (p = .53). Following outlier removal, there was a non-significant three-way interaction between OT/OTA, sex, and LPS/saline on the number of sweeps (b = -.340 ± .198, p = .055, F(1,40) = 3.88, conditional R^2^ = .64, marginal R^2^ = .38) (Supplemental Figure 12.D). Among males, there was no OT (p = .262) or OTA (p = .690) effect on sweep number relative to control after LPS (Supplemental Figure 12.D). In comparison, OTA females had a trending decrease in number of sweeps following LPS (b = -.271 ± .138, p = .055, F(1,40) = 3.88, conditional R^2^ = .702, marginal R^2^ = .38) relative to control females, with no difference in OT females (p = .107) and no effect of behavioural phenotype (p = .52) (Supplemental Figure 12.D). The average principal frequency of calls increased following LPS (b = 5.10 ± 1.60, p = .003, F(1,41) = 10.13, marginal R^2^ = .45), and control huddlers did not increase average call frequency following LPS-treatment (b = −9.45 ± 3.34, p = .008, F(1,41) = 7.89, conditional R^2^ = .59, marginal R^2^ = .52) (Supplemental Figure 12.E). The average principal frequency of female calls was higher than that of males (b = 3.64 ± 1.61, p = .029, F(1,40) = 5.14, marginal R^2^ = .45) (Supplemental Figure 15.G). Sinuosity increased following LPS (b = .36 ± .14, p = .015, F(1,41) = 6.39, conditional R^2^ = .24, marginal R^2^ = .16), suggesting an increase in frequency modulation (Supplemental Figure 12.F). There was no difference in sinuosity across treatment, behavioural phenotype, or sex (p = .54) (Supplemental Figure 15.H).

**Supplemental Figure 12.**
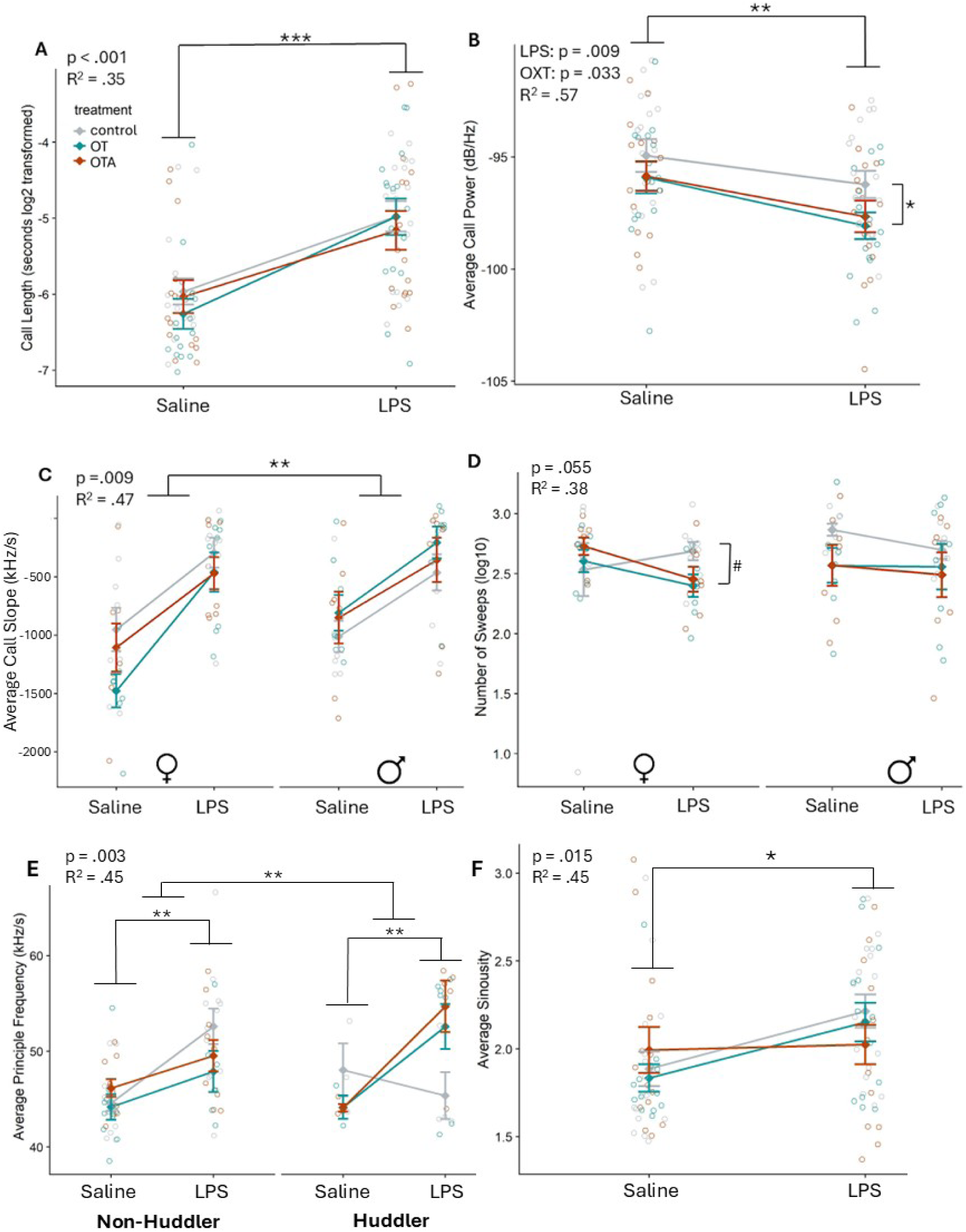
Following LPS treatment, average call length, slope, principle frequency, and sinuosity increased, while average call power decreased. A) Log2 transformed sweep call length (s) increased following LPS relative to saline (p < .001, conditional R^2^ = .46, marginal R^2^ = .35). B) Mice vocalized with reduced relative power (dB/Hz) in response to both LPS relative to saline (p = .009, conditional R^2^ = .77, marginal R^2^ = .57) and OT relative to controls (p = .033, conditional R^2^ = .77, marginal R^2^ = .57). C) Call slope (kHz/s) increased, becoming less negative, following LPS (p < .001, conditional R^2^ = .49, marginal R^2^ = .47) Additionally, there was an interaction effect of call slope with OT and sex, such that females treated with OT had more negatively sloped calls (p = .009, conditional R^2^ = .49, marginal R^2^ = .47). D) There was a non-significant three-way interaction between OT/OTA, sex, and LPS/saline on the number of sweeps (p = .055, conditional R^2^ = .64, marginal R^2^ = .38), such that OTA treated females trended towards having a reduced number of sweeps following LPS. E) The average principal frequency of calls increased following LPS relative to saline (p = .003, marginal R^2^ = .45) but control huddlers did not increase average call frequency following LPS-treatment (p = .008, conditional R^2^ = .59, marginal R^2^ = .52) . F) Sinuosity increased following LPS (p = .015, conditional R^2^ = .24, marginal R^2^ = .16).

### (h) Olfactory investigation positively correlated with average vocalization frequency and length after LPS

We ran a linear regression model including number of olfactory investigations, sex, and LPS/saline treatment on average call length and principal frequency. There was an interaction effect with LPS and focal mouse olfactory investigation on average call length and frequency, revealing a positive relationship between olfactory investigation (sniffs) and call length (b = .011 ± .004, p = .013, F(1,70) = 6.5, marginal R^2^ = .36) and frequency (b = 2.79 ± 1.00, p = .008, F(1,68) = 7.5, marginal R^2^ = .49) after LPS, but not after saline (Supplemental Figure 13). There was no main effect of LPS (p = .147), OT (p = .51), or OTA (p = .64) on olfactory investigation behaviour in the focal mouse and no sex effect on call length (p =.29) or frequency (p = .09) (Supplemental Figure 15.I, J)

**Supplemental Figure 13.**
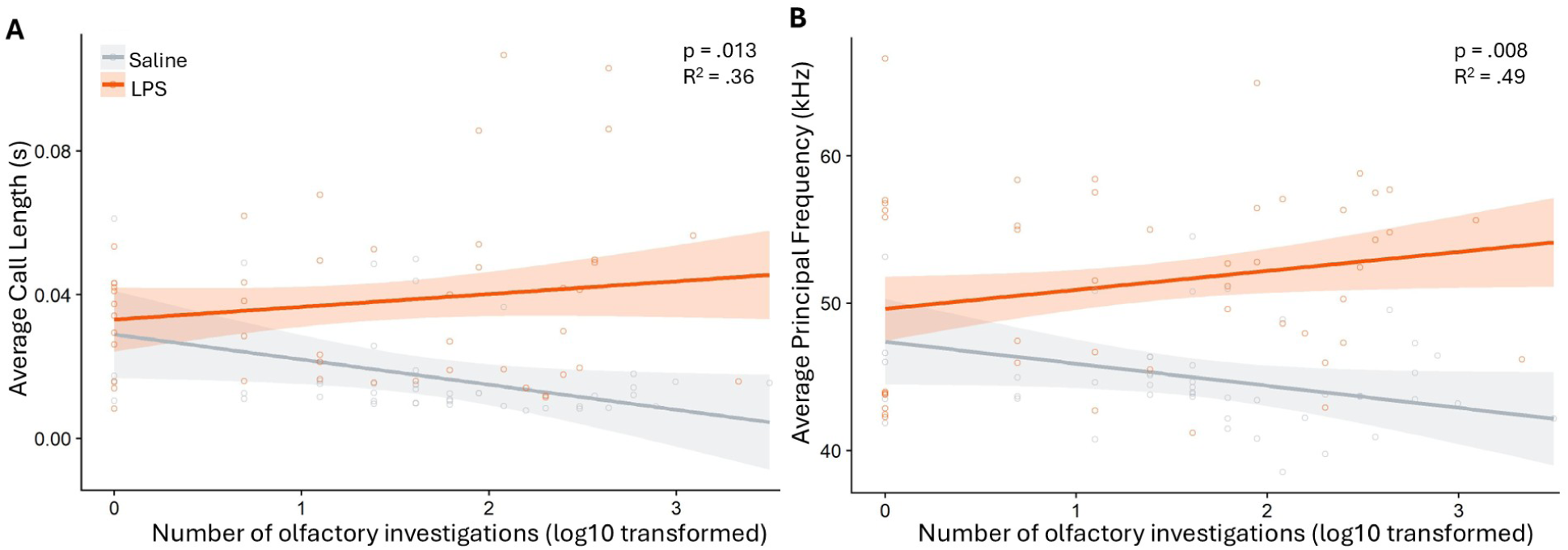
There was an interaction between olfactory investigation (number of sniffs), and LPS on critical call features: call length and principal frequency. After LPS, the positive correlation increased between (A) olfactory investigation and call length (p = .013, marginal R^2^ = .36) and (B) between olfactory investigation and average principal frequency (p = .008, marginal R^2^ = .49).

### (i) Individual data points for changes in call length and spectrogram

**Supplemental Figure 14.**
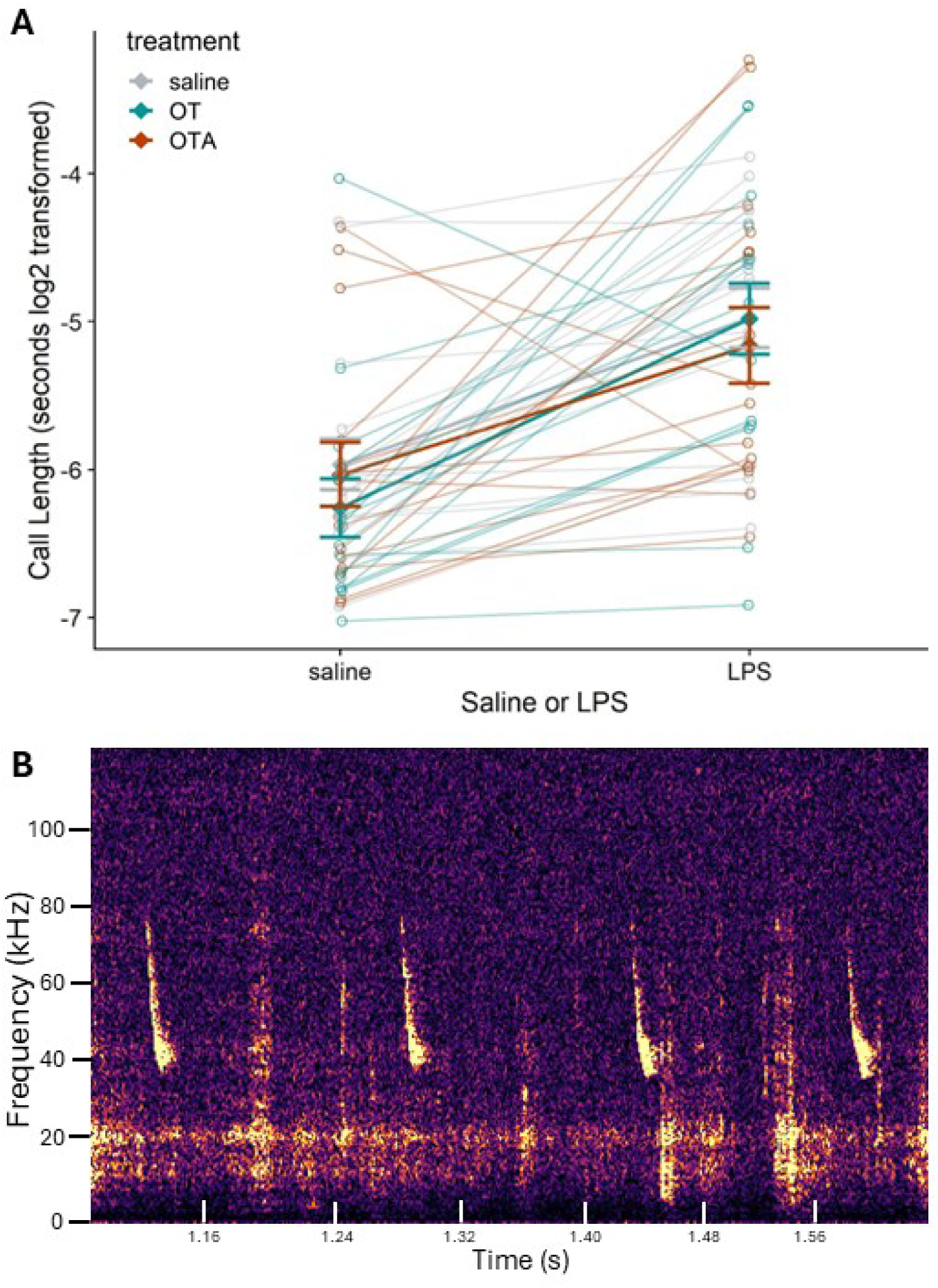
A) Change in sweep call length from saline to LPS with individual data points connected to demonstrate how individual mice changed in average call length over time. B) Spectrogram image of sweep vocalizations (Avisoft Bioacoustics, Berlin, Germany). Frequency (kHz) is shown on the y-axis, and minutes passed during recording is shown on the x-axis.

### (j) Figures with breakdown by sex

Here we include alternative figures to the main figures with a breakdown by sex when sex was omitted from the main figure for interpretability, but sex was included in the statistical analysis.

**Supplemental Figure 15.**
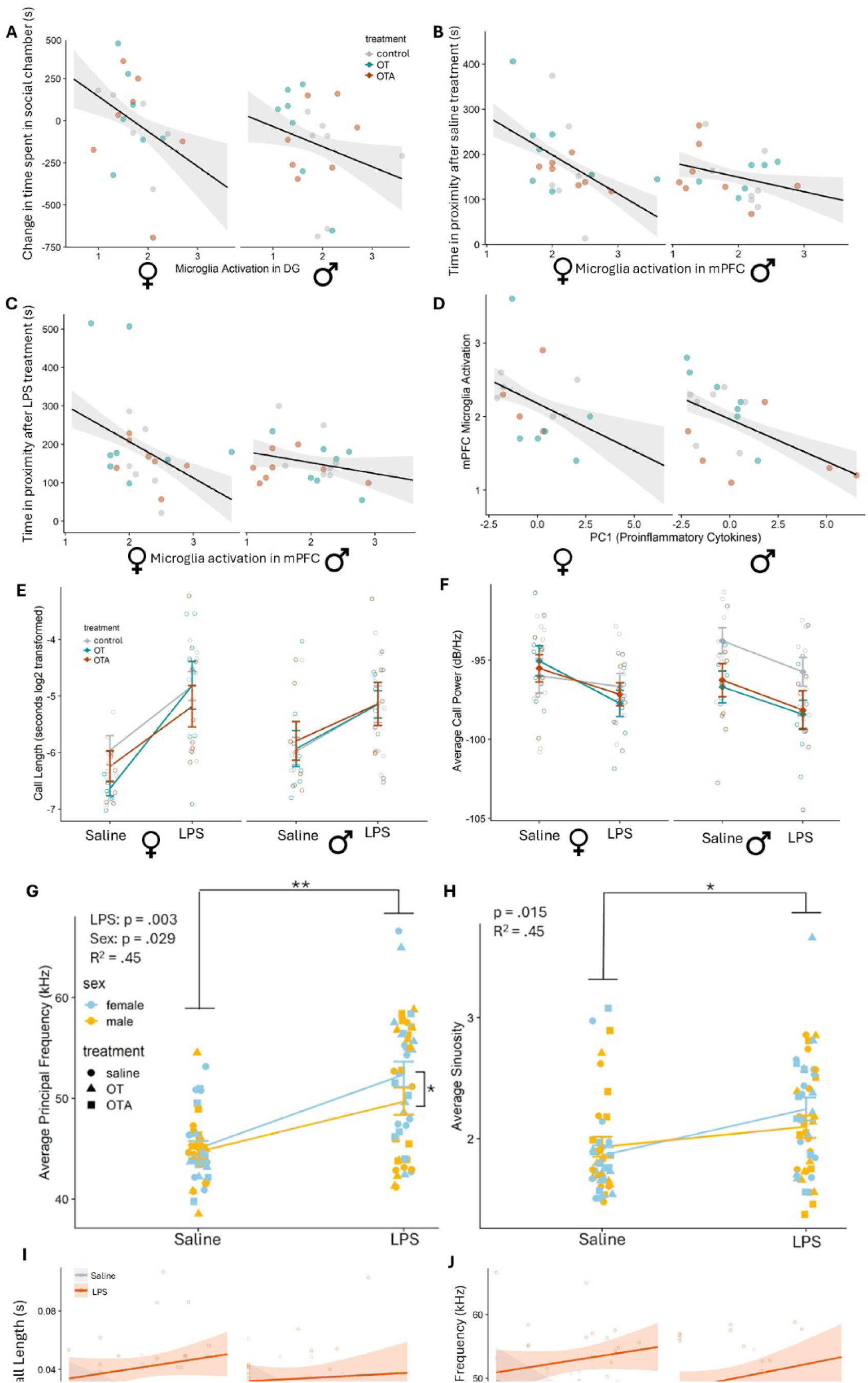
Extra figures that correspond to figures in the main text, but divided by sex. Because no sex differences were found in these analyses, the main figures do not show sex for ease of interpretability of main effects. Stats for significant effects are available in the main text. A) Corresponds to Figure 3.A. There was no sex interaction effect of microglia activation in the dorsal DG and sociability (p = .635). B) Corresponds to Figure 4.A. There was no sex difference with regards to microglial activation in the mPFC and time in proximity following saline injection (p = .091). C) Corresponds to Figure 4.B. There was no sex difference with regards to microglial activation in the mPFC and time in proximity following LPS injection (p = .124). D) Corresponds to Figure 5.A. There was no difference between sexes on the relationship between PC1 and mPFC microglial activation (p = .180). E) Corresponds to Supplemental Figure 12.A. There was no effect of sex on call length (p = .35). F) Corresponds to Supplemental Figure 12.B. There was no effect of sex on average call power (p = .56). G) Corresponds to Supplemental Figure 12.C. The average principal frequency of female calls was higher than that of males (b = 3.64 ± 1.61, p = .029, F(1,40) = 5.14, marginal R^2^ = .45). I) Corresponds to Supplemental Figure 13.A. There was no sex effect on call length (p =.29). J) Corresponds to Supplemental Figure 13.B. There was no sex effect on call frequency (p = .09).

